# The soluble guanylyl cyclase pathway is inhibited to evade androgen deprivation-induced senescence and enable progression to castration resistance

**DOI:** 10.1101/2023.05.03.537252

**Authors:** Ling Zhang, Clara I. Troccoli, Beatriz Mateo-Victoriano, Laura Misiara Lincheta, Erin Jackson, Ping Shu, Trisha Plastini, Wensi Tao, Deukwoo Kwon, Xi Chen, Janaki Sharma, Merce Jorda, James L. Gulley, Marijo Bilusic, Albert Craig Lockhart, Annie Beuve, Priyamvada Rai

**Author notes:** To whom correspondence should be addressed: Priyamvada Rai Papanicolau Building 131, University of Miami Miller School of Medicine 1550 NW 10^th^ Ave, Miami, FL 33136. These authors contributed equally to this study (Ling Zhang, Clara I. Troccoli).

## Abstract

Castration-resistant prostate cancer (CRPC) is fatal and therapeutically under-served. We describe a novel CRPC-restraining role for the vasodilatory soluble guanylyl cyclase (sGC) pathway. We discovered that sGC subunits are dysregulated during CRPC progression and its catalytic product, cyclic GMP (cGMP), is lowered in CRPC patients. Abrogating sGC heterodimer formation in castration-sensitive prostate cancer (CSPC) cells inhibited androgen deprivation (AD)-induced senescence, and promoted castration-resistant tumor growth. We found sGC is oxidatively inactivated in CRPC. Paradoxically, AD restored sGC activity in CRPC cells through redox-protective responses evoked to protect against AD-induced oxidative stress. sGC stimulation via its FDA-approved agonist, riociguat, inhibited castration-resistant growth, and the anti-tumor response correlated with elevated cGMP, indicating on-target sGC activity. Consistent with known sGC function, riociguat improved tumor oxygenation, decreasing the PC stem cell marker, CD44, and enhancing radiation-induced tumor suppression. Our studies thus provide the first evidence for therapeutically targeting sGC via riociguat to treat CRPC.

**Statement of significance:** Prostate cancer is the second highest cancer-related cause of death for American men. Once patients progress to castration-resistant prostate cancer, the incurable and fatal stage, there are few viable treatment options available. Here we identify and characterize a new and clinically actionable target, the soluble guanylyl cyclase complex, in castration-resistant prostate cancer. Notably we find that repurposing the FDA-approved and safely tolerated sGC agonist, riociguat, decreases castration-resistant tumor growth and re-sensitizes these tumors to radiation therapy. Thus our study provides both new biology regarding the origins of castration resistance as well as a new and viable treatment option.

## Introduction

Prostate cancer (PC) is the second leading cause of cancer-related death among American men. Despite an initially favorable outcome under androgen deprivation (AD), PC inevitably recurs as castration-resistant prostate cancer (CRPC)^1^ which is fatal and therapeutically underserved. To eradicate incurable PC, new therapies are critically needed, and an incomplete understanding of the molecular mechanisms underlying CRPC etiology is a barrier to therapeutic innovation.

We and others previously reported that AD induces senescence in castration-sensitive prostate cancer (CSPC) cells and tumors, as a tumor suppression mechanism^2–6^. We further reported that evasion of AD-induced senescence (ADIS) promotes outgrowth of emergent CRPC variants^4^. Indeed, we developed an early CRPC model from parental CSPC LNCaP cells through enrichment of a cell subpopulation that does not undergo an irreversible growth arrest in response to AD^4^. These ADIS-resistant counterparts (denoted as SB5) possess several traits of castration resistance including the ability to proliferate and retain androgen receptor (AR) expression under AD and to form tumors in castrated animals, albeit to a lesser extent than their fully CRPC counterpart LNAI cells^4, 7^. We then carried out unbiased transcriptomic analyses comparing the emergent CRPC LNCaP SB5 cells to their parental CSPC line (denoted as SB0) to identify new targets implicated in the progression to CRPC. We recently published a study validating one such target, thioredoxin-1 (TRX1); TRX1 is elevated in CRPC versus CSPC as a redox-protective response to AD-induced reactive oxygen species (ROS), and its genetic or pharmacologic inhibition re-sensitizes CRPC cells to AD^7^.

We also uncovered targets that were significantly decreased in the early androgen-deprived CRPC cells compared to their CSPC counterpart. These are likely candidates for mechanisms that serve to restrain CRPC development. One such target was the soluble guanylyl cyclase (sGC) complex. When activated by freely diffused NO, the sGC complex catalyzes the conversion of 5’-guanosine triphosphate (5’-GTP) to its second messenger molecule, cyclic guanosine monophosphate (cGMP)^8^. Through cGMP, sGC exerts anti-inflammatory, anti-proliferative and vasodilatory functions via downstream effectors, cGMP-dependent kinases (PKGs), cGMP-regulated phosphodiesterases (PDEs), and cyclic nucleotide-gated ion channels (CNEs)^9^.

Differential expression of its obligate heterodimeric subunits, sGCα1 and sGCβ1, as well as altered sGC activity has been previously reported in a panel of cancer cell lines^10^. Reduced/altered expression of either or both of the sGC subunits is associated with breast cancer progression^11–13^ and malignant gliomas^14^. These findings suggest a tumor-inhibitory function for sGC signaling. However, mechanistic roles for sGC involvement in cancer progression in general, or in prostate cancer progression, are not well-understood. This is a significant omission considering that CRPC is therapeutically under-served. Furthermore, stimulating the sGC pathway via clinically approved on-target agonists is a safe and well-tolerated intervention with minimal side-effects for pulmonary hypertension^9, 15^. Therefore, in this study, we investigated whether inhibition of sGC complex activity permits CRPC emergence, and whether an FDA-approved sGC agonist limits CRPC growth.

## Results

### Progression to castration resistance is associated with dysregulated sGC heterodimer expression and attenuated sGC activity

To study the molecular changes associated with resistance to androgen deprivation, we had previously developed an isogenic in vitro CRPC progression model from castration-sensitive prostate cancer LNCaP cells. Specifically, cyclical androgen deprivation of LNCaP SB0 cells promoted the outgrowth of counterpart AD-refractory LNCaP SB5 clones^4^. The SB5 cells resisted ADIS to proliferate in androgen-deprived culture media via charcoal-stripped serum (CSS)^4^ and could form tumors in castrated, male athymic mice^7^, thus confirming it is an early CRPC variant of the CSPC LNCaP SB0 parental line. We identified the sGC pathway (**Fig. 1A**) as an example of a CRPC-restraining mechanism from our unbiased screening of these emergent LNCaP-derived CRPC SB5 cells^4^. Analyses of differentially expressed genes from the transformed and normalized Illumina datasets, comparing the transcriptomes of proliferating and senescent LNCaP SB0 CSPC cells as well as those of LNCaP SB0 and their emergent CRPC LNCaP SB5 counterparts revealed that the gene expression signature for NO-sGC-cGMP signaling was enhanced in the senescent LNCaP SB0 vs. their proliferating counterparts (ADIS induction, **Fig. 1B**) and decreased in the emergent CRPC LNCaP SB5 subpopulations under AD (ADIS evasion, **Fig. 1B**).

**Figure 1.**
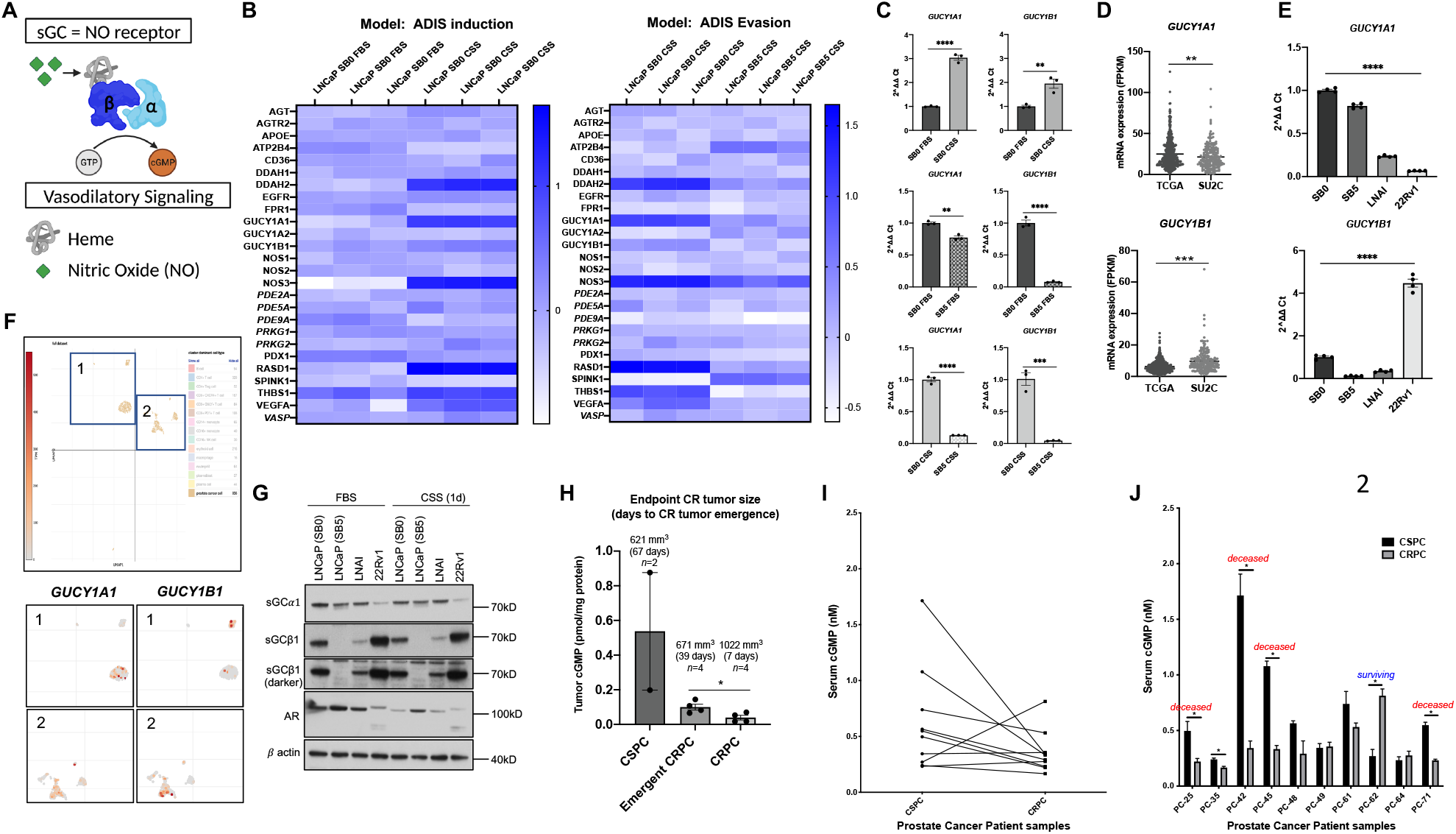
Decreased sGC complex activity correlates with preclinical as well as clinical progression to castration resistance. Note that all error bars represent ± SEM and * p< 0.05, ** p≤ 0.01, *** p≤ 0.001, **** p≤ 0.0001. A. The α1β1 heterodimer is the predominant form of the soluble guanylyl cyclase (sGC) complex, which binds nitric oxide (NO) to catalytically generate cGMP, which mediates downstream vasodilatory phenomena. B. Heat maps of genes differentially altered in LNCaP SB0 cultured in FBS- vs. CSS-supplemented media (denoted as ‘ADIS Induction’) and LNCaP SB5 vs LNCaP SB0 cultured in CSS-supplemented media (denoted as ‘ADIS Evasion’). Increasing blue intensity signifies increased expression. These analyses indicate the NO-sGC-cGMP signaling reactome is enhanced in the CSPC LNCaP SB0 cells which undergo ADIS, and is downregulated in the emergent CRPC variants, LNCaP SB5, which can proliferate under AD unlike their SB0 counterparts. C. The mRNA levels for *GUCY1A1* (α1 subunit gene) and *GUCY1B1* (β1 subunit gene) are altered in opposing directions for CSPC SB0 vs. emergent CRPC SB5 cells under AD. Levels were measured by qPCR for the indicated cells and conditions. *ActinB* was used as the normalization control. Cells were cultured for 5 days in CSS-supplemented media to induce AD. The p-values were established via an unpaired two-tailed Student’s t test. D. Analysis of mRNA levels for *GUCY1A1* and *GUCY1B1* in prostate tumor samples derived from the TCGA (largely non-metastatic PC) and SU2C (metastatic) public datasets show differences in expression between the two sGC subunits. Transcript levels are expressed as fragments per kilobase of transcript per million (FPKM), and p-values are from an unpaired two-tailed Student’s t test. E. *GUCY1A1* and *GUCY1B1* mRNA levels across a panel of CSPC and CRPC cell lines (LNCaP SB0, LNCaP SB5, LNAI, 22Rv1) were analyzed by qPCR and indicate stoichiometric dysregulation of the heterodimer. *ActinB* was used for normalization. Significance was assessed by an unpaired two-tailed Student’s t test. F. Metastatic treatment-resistant PC tumors exhibit non-stoichiometric *GUCY1A1* and *GUCY1B1* expression within different tumor regions. UMAP plots from publicly available scRNA-seq data^19^ are shown (top: only the PC cells are highlighted, middle, bottom: specific sections of the tumor corresponding to the numbered regions (top) are expanded to show the lack of spatial correlation between *GUCY1A1-* or *GUCY1B1-*expressing PC cells. G. Dysregulation of sGC subunits is seen at the protein level in CRPC cells. Immunoblotting of total protein lysates (15 µg) from LNCaP SB0, SB5, LNAI and 22Rv1 cells (FBS-supplemented culture) against sGC⍺1 and sGCβ1 with β actin as the loading control. Molecular weights of markers run on the original immunoblot are indicated on the right. H. Lower intratumoral cGMP levels correlate with faster CRPC tumor growth. Intratumoral cGMP levels were measured in flash-frozen tissue from subcutaneous xenograft tumors derived from CSPC LNCaP SB0 cells (n=2), emergent CRPC LNCaP SB5-derived cells (n=4) and fully CRPC LNAI cells (n=4). Endpoint tumor volumes at time of harvest, number of distinct tumors evaluated, and time to palpable CR tumor formation are shown for each category. Significance was assessed by an unpaired two-tailed Student’s t test. I. Trends for sera cGMP levels, pre- vs. post-castration resistance, are shown for matched CSPC/CRPC samples from 10 de-identified PC patients. J. The CSPC vs. CRPC sera cGMP levels from the 10 patients in (I) are shown. Note that patients 25, 42, 45, 71 are deceased; patient 62 had stable disease at comparable time of assessment (see Fig. S1H). Significance was assessed by an unpaired two-tailed Student’s t test.

We chose to focus on the sGC component of this interactome as it was in the top 0.01% of all significantly altered genes in our dataset, and is therapeutically actionable and understudied in PC. Independent verification of the transcriptomics data via qPCR analysis of sGCα1 (*GUCY1A1*) and its obligate stoichiometric heterodimeric partner sGCβ1 (*GUCY1B1*) was carried out in the proliferating vs. senescent SB0 as well as SB5 vs. SB0 counterparts. Consistent with our unbiased profiling, we found both subunits were elevated in senescent SB0 cells vs. their proliferating counterparts (**Fig. 1C,** top). By contrast, both sGC subunits were significantly downregulated in the ADIS-resistant SB5 cells (especially under AD culture), relative to the parental CSPC SB0 cells (**Fig. 1C**, middle, bottom). We also evaluated mRNA levels of both sGC heterodimer subunits in the CSPC LAPC4 SB0 cell line and its ADIS-resistant counterpart LAPC4 SB1^4^. We found that both subunit levels were also significantly lower in the ADIS-resistant SB1 variants versus their parental CSPC SB0 counterparts (**Supplementary Fig. S1A**).

Increases in cancer incidence or progression have not been noted in long-term clinical use of sGC agonists. Furthermore, unbiased CRISPR and RNAi-based gene dependency screens from <u>depmap.org</u> indicate that cancer cells (including PC) do not depend on either sGC subunit for their viability (**Supplementary Fig S1B, C).** These data are consistent with a non-essential role for sGC in cancer, suggesting its inhibition is unlikely to offer therapeutic benefits. Our data suggest the sGC complex is attenuated with PC progression. Therefore, we analyzed mRNA levels of the sGC α1 and β1 subunits, *GUCY1A1* and *GUCY1B1* respectively, in the TCGA (mainly CSPC) vs. SU2C (mainly metastatic) datasets in cBioportal (cbioportal.org) (**Fig. 1D**) as well as in primary vs. metastatic prostate cancers in the Grasso human PC dataset, which identified potential CRPC drivers^16^ (**Supplementary Fig. S1D**). Both showed that sGCα1 mRNA (*GUCY1A1)* was significantly downregulated in metastatic PC vs. primary/CSPC indicating dysregulation of the functional heterodimeric complex^17^. By contrast, the sGCβ1 *(GUCY1B1*) subunit was not as consistently altered across this continuum, with transcript levels being significantly higher in the SU2C cohort but significantly lower in the Grasso cohort (**Fig. 1D, Supplementary Fig. S1D**).

We also assessed the relative mRNA levels of these subunits in LNCaP SB0 (CSPC), LNCaP SB5 (emergent CRPC), their established CRPC counterpart, LNAI (all of which express full-length AR), and in the AR-V7-expressing CRPC line, 22Rv1. We found that *GUCY1A1* levels were consistently lower in the CRPC lines relative to the CSPC LNCaP line (**Fig. 1E)**. However, while *GUCY1B1* was also lower in the AR full-length CRPC lines, it was significantly elevated in the 22Rv1 line. Indeed, 22Rv1 also exhibited stoichiometric dysregulation of the sGC complex, with very low *GUCY1A1* in conjunction with very high *GUCY1B1* (**Fig. 1E).** Analysis of *GUCY1A1, GUCY1B1* levels from CSPC, CRPC and NEPC cell lines (depmap.org) support our findings that the sGC complex is attenuated during PC progression, with the NCI H660 NEPC line exhibiting no detectable expression of either subunit (**Supplementary Fig S1E**). Analysis of sGC subunit expression in single-cell RNA-seq datasets of metastatic prostate cancer^18^ indicated they are expressed mainly in the cancer cells (**Supplementary Fig. S1F**). However, the areas of *GUCY1A1* and *GUCY1B1* expression in the prostate cancer are not coincident (**Fig. 1F, Supplementary Fig. S1F**), suggesting the areas of tumor sampling for transcriptomics can affect the levels of the two subunits in bulk sequencing datasets. Thus, the expression-level dysregulation between the sGC subunits observed in our emergent CRPC model (**Fig. 1C**) reflected the spatially heterogenous expression of these subunits seen in tumor single-cell transcriptomics, rather than the bulk transcriptomics from patient datasets. At the protein level (**Fig. 1G**), sGCβ1 better matched its mRNA levels across the cell lines relative to sGCα1 protein, which remained largely unchanged across the AR-FL lines but was low in 22Rv1, similar to its mRNA levels (**Fig. 1E**). Acute androgen deprivation (reflected in decreased AR-FL levels) did not significantly affect protein levels of either subunit (**Fig. 1G**). The lack of consistent correlation between the transcriptional and protein levels of sGC complex subunits we note here has previously been reported for some cancer cell lines, including PCa lines (Fig 1 vs. 2, therein)^10^. This inconsistency may reflect relative differences in β1 subunit vs. α1 subunit protein stability in PC cells.

The physiologic activities of the sGC complex arise through cGMP signaling, and because phosphodiesterase 5A (PDE5A) levels are low in prostate cancer^19^ and especially so in advanced stages **(Supplementary Fig. S1G**), we reasoned cGMP levels in PC mainly reflect sGC activity. Therefore, we evaluated the effects of dysregulated heterodimer expression on sGC activity. The patient datasets we analyzed in Fig. 1D, Supplementary Fig. S1D represent established CRPC tumors. We wanted to evaluate whether sGC signaling declines in cells that progressively acquire castration resistance, to recapitulate the setting in which we identified the involvement of this pathway in PC progression. We therefore measured cGMP levels in castrate-resistant (CR) tumors formed by the parental CSPC LNCaP SB0 line, the emergent CRPC LNCaP SB5 variant and the established CRPC LNCaP variant, LNAI. As anticipated, following injection of cells into castrated male immunocompromised mice, these lines respectively produced CR tumors of similar size at progressively shorter durations (**Fig. 1H**). Moreover, tumors from cell lines with a greater capacity for CR growth exhibited correspondingly lower cGMP levels (**Fig. 1H**), reinforcing the idea that attenuation of sGC signaling occurs as cells progress to castration resistance.

Finally, we measured cGMP levels, in a disease outcome-blinded fashion, in matched sera taken before and after progression to CRPC from 10 prostate cancer patients. The overall trends supported that lower cGMP was associated with the castration-resistant state (**Fig. 1I**). We next evaluated the observed trends of cGMP alteration in CRPC vs. CSPC against clinical outcomes in these patients (**Supplementary Fig. S1H**). Significant declines in cGMP levels with CRPC progression (patients 25, 42, 45, 71) correlated with rapid disease progression and mortality following CRPC diagnosis. By contrast, patient 62, who showed an increase in cGMP following castration resistance, had less aggressive disease and prolonged survival post diagnosis (**Fig. 1J, Supplementary Fig. S1H**). With the obvious caveat of the small numbers, our results here suggest declining cGMP levels could accurately predict disease progression and fatality in a random patient dataset, through non-invasive sera cGMP profiling. These data thus support our observations in cell lines and xenograft tumor models that attenuated sGC signaling is associated with progression to CRPC.

### Stoichiometric dysregulation of sGC subunits and inhibition of sGC activity are early events in CRPC progression

The heterodimeric structure of the sGC complex is essential to its catalytic activity^20^. Reduced or altered expression of either subunit of the sGC complex leads to impaired downstream signaling, and is implicated in several pathologies, including cardiovascular^21–23^, neurodegenerative^24^ and genitourinary disorders^25^. However, whether this phenomenon plays a role in cancer is unknown, to the best of our knowledge. We particularly wished to determine if stoichiometric dysregulation occurs as an early event in the progression to CRPC, as suggested by our transcriptomics profiling. Therefore, using a panel of isogenic lines representing early-stage but progressively more castration-resistant counterparts, we assessed expression levels of the heterodimer subunits as well as the ability of the sGC complex to be functionally stimulated. These LNCaP counterparts were established in our lab and are denoted as follows **(Supplementary Fig. S2A**): SB0 (parental CSPC), SB5 (emergent CRPC), SB5X (derived from a castration-resistant tumor formed by SB5), SB5XX (derived from a castration-resistant tumor formed by the SB5X line). The SB5X line formed CR tumors more readily in vivo relative its counterpart SB5 line (**Supplementary Fig. S2B, C**) just as the SB5 cells formed CR tumors more readily than the parental SB0 line^7^.

Analyses of sGC subunit expression in this emergent CRPC panel indicated stochiometric dysregulation occurred most dramatically for the obligate heterodimeric β1 subunit, which is significantly downregulated at both the mRNA and protein level in SB5, SB5X, SB5XX lines (**Fig. 2A, B**). Differing from our advanced PC patient dataset analyses in **Fig. 1**, the α1 subunit was downregulated less consistently than the β1 subunit at the message level in these emergent CR lines (**Fig. 2A**). This held particularly true under androgen-deprived (charcoal stripped serum; CSS) conditions versus the androgen replete (fetal bovine serum; FBS). The findings under AD are particularly striking as they reflect 30 hours of CSS culture, a relatively acute duration that does not produce major decreases in AR or chronic effects such as senescence or other anti-tumor responses^4^. This suggests that the effects on the β1 subunit are unlikely to be AR-regulated. At the protein level, similar to what we observed in **Fig. 1G**, there were insignificant differences in sGCα1 expression across these progressively CR cell lines but there was a striking loss of the sGCβ1 protein (**Fig. 2B**).

**Figure 2.**
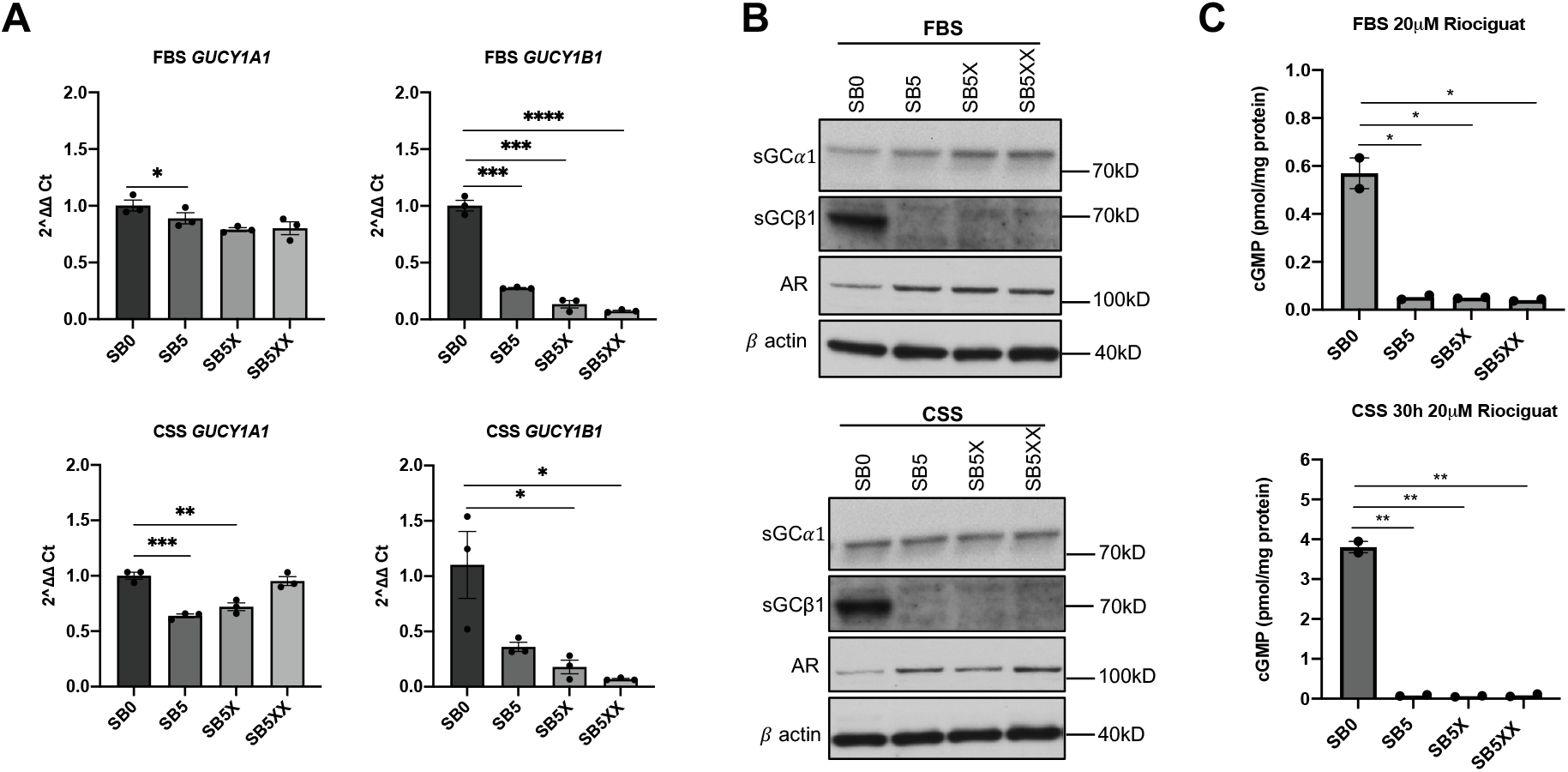
Emergent CRPC variants derived from parental LNCaP CSPC cells exhibit significant loss of sGCβ1 expression and activity. Note that all error bars represent ± SEM and * p< 0.05, ** p≤ 0.01, *** p≤ 0.001, **** p≤ 0.0001. A. The mRNA levels of *GUCY1A1* and *GUCY1B1* were measured by qPCR in our panel of isogenically matched cell variants of progressive castration-resistance: LNCaP SB0 (parental CSPC), and the counterpart emergent CRPC lines, SB5, SB5X, and SB5XX under the indicated culture conditions. *ActinB* was used as the normalization control. The p-values are from an unpaired two-tailed Student’s t test. B. Western blotting of total protein lysates (15 µg) from the lines in (A) indicate loss of sGCβ1 protein in emergent CRPC lines vs. their CSPC SB0 counterpart. Immunoblotting against both sGC subunits as well as AR expression, with β actin as the loading control is shown. To assess effects of acute androgen deprivation, cells were cultured for 30 hours in CSS media. Molecular weights of the markers from the original immunoblot are indicated on the right. C. Emergent CRPC cells show significantly dampened sGC activity. The cell lines indicated in (A) were cultured in FBS or CSS media for 30 hours and then treated with 20 µM riociguat for 30 minutes, to stimulate sGC signaling, following which cGMP levels were measured. The p-values are from an unpaired two-tailed Student’s t test.

Nevertheless, β1 subunit downregulation was sufficient to essentially suppress sGC signaling, as seen by the inability of the FDA-approved sGC agonist, riociguat^26^ (aka Adempas, BAY 63-2521), to stimulate cGMP production in any of the emergent CRPC cell lines relative to their CSPC parental SB0 counterpart (**Fig. 2C**). Thus, collectively these data support that sGC stoichiometric dysregulation is an early occurrence in the progression to CRPC. This is associated with drastic transcriptional loss of the obligate β1 sGC heterodimeric subunit, leading to abrogation of sGC catalytic activity and cGMP production.

### Inhibition of sGC signaling via sGCβ1 knockout facilitates evasion of androgen deprivation-induced senescence (ADIS) and castration resistant growth

We next sought to determine if the loss of sGC activation has a functional role in the acquisition of castration resistance in CSPC cells. Gene set enrichment analysis (GSEA)^27^ of our transcriptomics data indicated that NO signaling genes are enriched in senescent LNCaP SB0 vs. their proliferating counterparts (**Supplementary Fig. S3A**). The CRPC progression lines that showed loss of sGC activation (**Fig. 2C**) also become increasingly resistant to ADIS, as seen by the progressively lower induction of p16^INK4a^, the molecular pathway underlying ADIS^4^ (**Fig. 3A**). Thus, we postulated that the NO-sGC-cGMP signaling enforces ADIS and that its abrogation is required to overcome the ADIS barrier to castration resistance.

**Figure 3.**
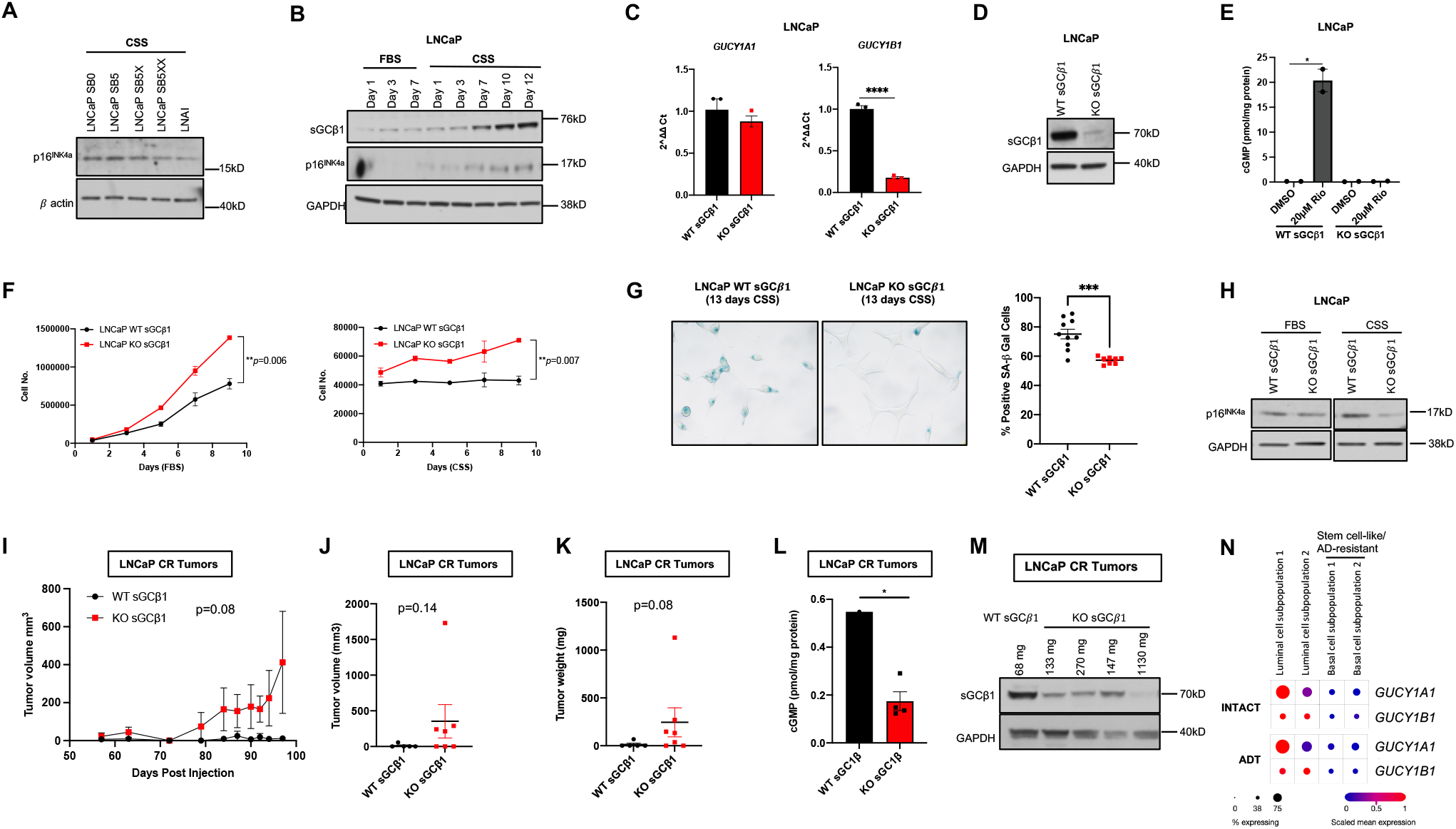
Downregulation of sGC signaling facilitates ADIS evasion and castration-resistant tumor growth in CSPC cells. Note that all immunoblots show the molecular weight of the markers run on the original membrane on the right. All error bars represent ± SEM and * p< 0.05, ** p≤ 0.01, *** p≤ 0.001, **** p≤ 0.0001. A. Emergent CRPC and fully CRPC panel of cell lines show progressively lower p16INK4a induction under AD culture compared to parental SB0 cells, indicating their ability to evade ADIS. Total protein lysates (15 µg) were immunoblotted against the indicated proteins. B. ADIS is associated with increasing levels of sGCβ1 protein. Total LNCaP SB0 protein lysates (15 µg) harvested at the specified time points and culture conditions were immunoblotted against the indicated proteins. C. The mRNA levels of *GUCY1A1* and *GUCY1B1* were measured via qPCR in LNCaP cells deleted for sGCβ1 by CRISPR-Cas9 modification (KO sGCβ1). *ActinB* was used for normalization. The p-values were determined by an unpaired two-tailed Student’s t test. D. Knockout of sGCβ1 in LNCaP cells was validated by immunoblotting for loss of protein. Total protein lysates (10 µg) for sGCβ1 levels from counterpart control and KO sGCβ1 LNCaP cell lines were immunoblotted. E. Knockout of sGCβ1 abrogates sGC activity. Baseline and riociguat-stimulated cGMP levels from control and counterpart KO sGCβ1 LNCaP lines are shown. The p-values were determined by an unpaired two-tailed Student’s t test. F. Knockout of sGCβ1 in LNCaP CSPC cells enables their proliferation under AD culture. Cell proliferation curves were established by plating 50,000 cells for KO sGCβ1 or matched control LNCaP cells and monitoring growth in FBS-supplemented (left) or CSS-supplemented (right) media over 9 days. The p values were determined by comparing WT sGCβ1 vs KO sGCβ1 cell numbers at day 9 using unpaired two-tailed Student’s t tests. G. Knockout of sGCβ1 decreases number of CSPC cells that undergo ADIS. WT and counterpart KO sGCβ1 LNCaP cells were cultured under AD conditions for 13 days and then assayed for the senescence-associated beta-galactosidase (SA-beta-gal) activity (left). A minimum of 150 cells across 4-5 high powered fields per sample was evaluated for positively stained cells. Percentage positivity of staining was determined by the number of SA-beta-gal-stained cells normalized to the total number of cells per field (right). Unpaired two-tailed t-test with Welch’s correction was used to determine p-values, and outliers were excluded via the ROUT method (Q=1%). H. KO sGCβ1 LNCaP cells show lower p16^INK4a^ protein levels relative to its wt counterpart cells following prolonged culture in CSS-supplemented culture to induce ADIS. Western blotting of total protein lysates (10µg) is shown for the indicated cell lines, following culture in FBS or CSS media for 13 days. I. Loss of sGCβ1 promotes castration-resistant tumor growth by LNCaP CSPC cells. The KO sGCβ1 or matched WT LNCaP cells were subcutaneously injected at a single site per 5-week old, Nu/Nu, castrated male mice. Tumor formation kinetics are shown. Wilcoxon two-sample test was used to determine the p-value. J. Endpoint tumor volumes (mm^3^) for the KO sGCβ1 and matched WT groups (from I) are shown. Each point represents the tumor from an individual mouse. Final tumor incidence: WT 1/5; KO sGCβ1 4/7. Wilcoxon two-sample test was used to determine the p-value. K. Endpoint tumor weights (mg) in the KO sGCβ1 and matched WT sGCβ1 groups were measured. Each point represents an individual tumor. Wilcoxon two-sample test was used to determine the p value. Note that absence of a tumor mass upon necropsy is indicated as zero. L. Intratumoral cGMP levels are lower in KO sGCβ1 tumors relative to wt counterparts. The p-values were established via an unpaired two-tailed Student’s t test. M. Immunoblotting for sGCβ1 levels in tumor lysates (30µg) corresponding to the tumors evaluated for cGMP in (L) is shown. Endpoint tumor weights are indicated for each individual tumor. N. Analysis of *GUCY1A1* and *GUCY1B1* mRNA levels from four distinct epithelial clusters (two basal, two luminal) derived from human prostate tissue. Expression levels from publicly available scRNA-seq data from the Single Cell Portal at Broad Institute

The sGCβ1 protein is notably absent in LNCaP SB5 variants (**Fig. 2B**). Moreover, in LNCaP CSPC cells, sGCβ1 protein is progressively elevated with increased duration of AD, coincident with p16^INK4a^ induction (**Fig. 3B**). Loss of the sGCβ1 subunit is reported to eliminate physiologic sGC enzymatic function in cells and tissues from germline-deleted mice^28^. Therefore, we wished to test whether loss of the sGCβ1 subunit in LNCaP CSPC cells would also inhibit sGC activation and confer ADIS resistance. To do so, we utilized a CRISPR sGCβ1 knockout LNCaP line (**Fig. 3C, D**), and confirmed that deletion of β1 abrogated sGC signaling, as seen by the inability of sGCβ1 KO cells to be stimulated by riociguat (**Fig 3E**). Thus, loss of the β1 subunit in LNCaP cells recapitulates functional sGC loss. Under androgen-replete conditions, the sGCβ1 KO cells displayed enhanced proliferation compared to their WT counterparts (**Fig 3F,** left), consistent with cell cycle-inhibitory effects previously reported for the β1 subunit^29^. More significantly, we found that the LNCaP SB0 β1 KO cell population behaved similarly to the LNCaP SB5 variants^4^: they were able to proliferate under androgen-deprived conditions (**Fig. 3F,** right), displayed overall lower levels of senescence-associated beta-galactosidase (SA-beta-gal) activity (**Fig. 3G**) and lower p16^INK4a^ under prolonged AD culture (**Fig. 3H**).

We next subcutaneously injected either the LNCaP sGCβ1 KO or their counterpart WT cells into the flanks of castrated Nu/Nu male animals at a single site per mouse, and monitored tumor growth. Neither cohort of injected animals exhibited any significant differences in weight or showed other ill-effects **(Supplementary Fig. S3B**). LNCaP CSPC cells are not expected to efficiently form tumors in castrated mice. Notably, the tumor kinetics between the WT and sGCβ1 KO tumors were markedly different with more CR growth by the latter (**Fig. 3I**). Differences in endpoint tumor volumes (**Fig. 3J**) and weights (**Fig. 3K**) were borderline non-significant due to the wide variability in CR tumor size formed by the sGCβ1 KO cells and likely our relatively small sample sizes. Nevertheless, only a single LNCaP WT sGC injection site formed a very small CR tumor whereas 4 out of 7 sites injected with the LNCaP sGCβ1 KO cells robustly formed CR tumors (**Figs. 3I-K, Supplementary Fig. S3C)**. During necropsy, a fifth sGCβ1 KO site was found to have formed a very small tumor; however, none of the other WT sites showed any sign of tumor formation upon necropsy.

Moreover, the intratumoral cGMP levels were also much higher in the one WT CR that formed compared to the four sGCβ1 KO CR tumors (**Fig. 3L, Supplementary Fig. 3D**). Furthermore, as our CRISPR cells were pooled as opposed to being single clones, we assessed whether there was variability in sGCβ1 expression among the castrate-resistant tumors that formed. Significantly we found that the largest sGCβ1 KO tumor had near undetectable sGCβ1 levels compared to the smaller tumors (**Fig. 3M**), again reinforcing that repression of the functional sGC complex promotes castration-resistant growth by CSPC cells.

To our knowledge, this is the first finding to indicate sGC signaling inhibits progression to castration resistance. Our results are consistent with the fact that basal cells in human prostatic tissue, which are naturally AD-resistant and most likely possess prostate regenerative properties, have barely detectable sGC subunit levels compared to luminal cells, which give rise to most prostate adenocarcinomas^30^ (**Fig. 3N**).

Collectively, these findings support that inhibition of sGC signaling via loss of sGCβ1 lowers the ADIS barrier to emergence of castration resistant sub-populations but is perhaps not sufficient to fully confer castration resistance. The observed abrogation of sGCβ1 expression and inability to stimulate sGC activity in emergent CRPC variants (**Fig. 2**) is consistent with this idea. Thus, loss of sGC signaling appears to be a key adaptation on the path to castration resistance.

### Stimulation of sGC signaling cooperates with androgen deprivation to induce cell death in established castration-resistant cells

The sGC agonist, riociguat is a safe, well-tolerated FDA-approved treatment for pulmonary hypertension^31, 32^ and can stimulate sGC even in the absence of nitric oxide^32^. Given that CRPC progression is associated with decreased sGC expression and signaling, and that established CRPC cells and human tumors have lower cGMP levels than CSPC counterparts (**Fig. 1H-J**), we predicted that riociguat treatment would enhance the attenuated sGC pathway thus producing CRPC-restraining responses. We then assessed whether established LNCaP-derived fully CRPC LNAI^7^ cells respond to riociguat. Unlike the early CRPC SB5 variants (**Fig. 2C**), LNAI could be stimulated by riociguat to produce cGMP under androgen-replete conditions; however, ability to stimulate cGMP in LNAI was significantly lower than in parental CSPC LNCaP SB0 cells (**Fig. 4A**), supporting our idea that the sGC pathway is dampened in the progression to CRPC. Androgen deprivation (CSS), however, significantly increased cGMP levels in LNAI cells relative to androgen replete (FBS) culture (**Fig. 4B**). Consistent with this finding, when we treated LNAI and 22Rv1 CRPC cells with riociguat under FBS conditions, we found they were relatively resistant to riociguat-induced cytotoxicity but showed a striking sensitization under androgen-deprived culture (**Fig. 4C; Supplementary Fig. S4A**). Sensitization to riociguat was also observed in LNAI cells when combined with the gold-standard AR inhibitor, enzalutamide (**Fig. 4D**). By contrast, the parental CSPC counterparts, LNCaP SB0, showed a significant loss of viability under riociguat treatment regardless of androgen-replete or deprived conditions (**Supplementary Fig. S4B, C),** reflecting their much higher cGMP levels under riociguat stimulation compared to LNAI **(Fig. 4A, B).** This is consistent with the robust expression of both sGC subunits in LNCaP SB0 compared to SB5 or LNAI (**Fig. 1G**). However, whereas riociguat induces cell death in LNCaP SB0 based on loss of cell number and morphology under AD conditions (**Supplementary Fig. S4C**), it evokes senescence (reminiscent of ADIS) under androgen-replete conditions as seen from the enhanced SA-beta-gal staining (**Supplementary Fig. S4D**). This observation is consistent with our observation that enhanced sGC activity (**Fig. 4A**) is associated with reinforcement of tumor suppressive barriers preventing CSPC progression to CRPC.

**Figure 4.**
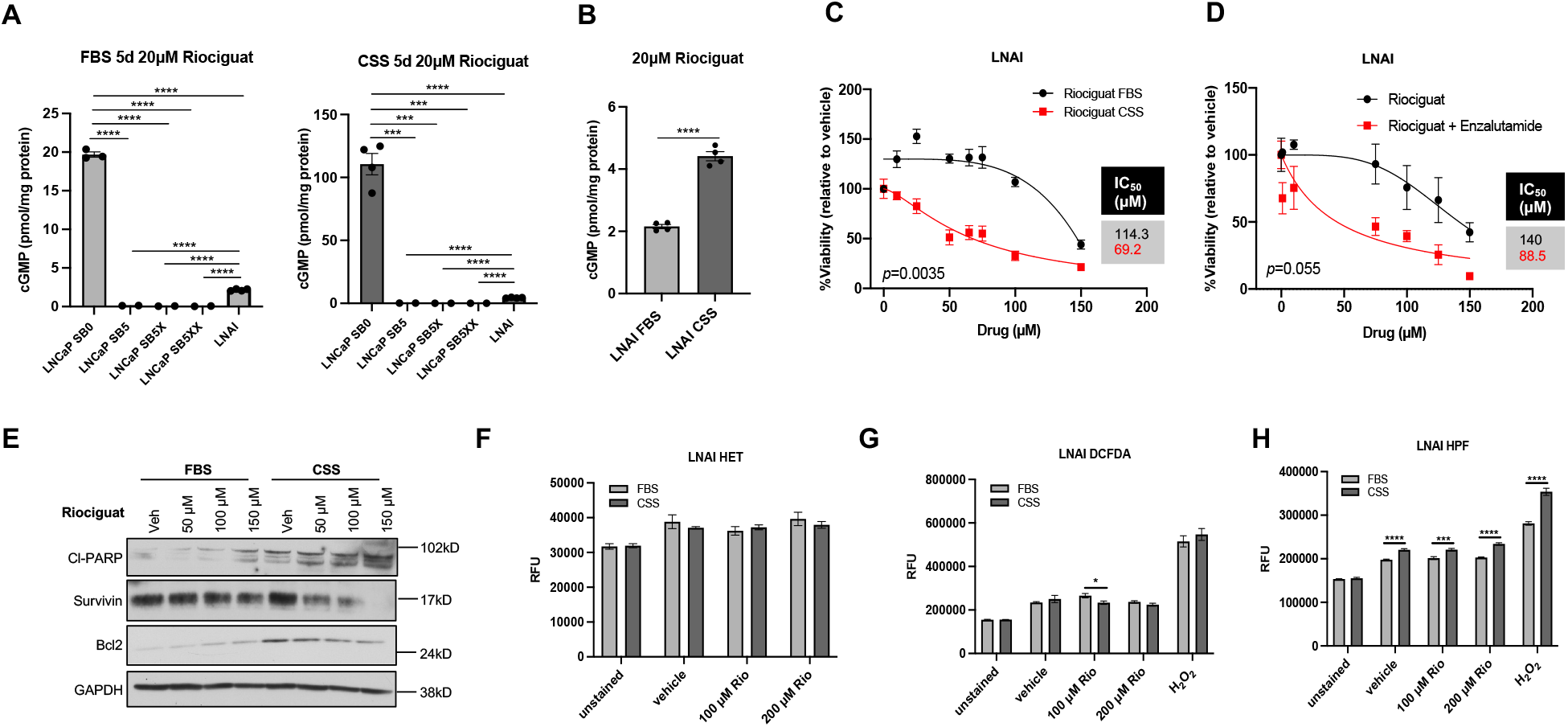
Stimulation of sGC signaling cooperates with androgen deprivation to induce apoptosis in CRPC cells. Note that all error bars represent ± SEM and * p< 0.05, ** p≤ 0.01, *** p≤ 0.001, **** p≤ 0.0001. A. sGC activity is dampened in established CRPC relative to CSPC, but elevated relative to emergent CRPC variants. LNCaP SB0, SB5, SB5X, SB5XX and LNAI cells were cultured in FBS- or CSS-supplemented media for 5 days, then treated with 20 µM riociguat for 30 minutes before measuring cGMP levels. p-values were determined by an unpaired two-tailed Student’s t-test. B. Comparison of relative sGC activity in androgen-replete (FBS) vs. androgen-deprived (CSS) LNAI CRPC cells shows AD enhances sGC activity. Values for cGMP levels from Fig.4A and Fig.4B are plotted together. p-values were determined by an unpaired two-tailed Student’s t-test. C. Androgen deprivation sensitizes CRPC cells to riociguat. LNAI cells were plated in triplicate in 96-well plates and treated with vehicle (DMSO) or riociguat in FBS or CSS media for 72 hours, with daily redosing. The IC_50_s for each curve were modeled using a four-parameter log-logistic function and are shown in the table beside each graph. Wilcoxon two-sample test was used to determine p-values. D. LNAI cells were plated in FBS media as in C, then incubated with 10 µM enzalutamide for 24 hours before dosing with the indicated various concentrations of riociguat alone or riociguat with enzalutamide for 72 hours, with daily redosing. Viability and IC_50_s as well as p-values were established as in Fig. 4C. Wilcoxon two-sample test was used to determine p-values. E. LNAI cells were plated in 10 cm dishes and treated with the indicated riociguat doses in either FBS or CSS media for 72 hours, with daily redosing. Following treatment, cells were harvested and immunoblotting of 26 µg total protein lysates was carried out against the specified antibodies. Molecular weights of the markers run on the original membrane are shown on the right. F. LNAI cells were plated in triplicate in 96-well plates and treated with vehicle (DMSO) or riociguat in FBS or CSS media for 20 hours. Intracellular superoxide levels were assessed by PI fluorescence, following incubation with 10 µM HEt. Unpaired two-tailed Student’s t-test were used to determine p-values. G. LNAI cells were plated in triplicate in 96-well plates and treated with vehicle (DMSO) or riociguat in FBS or CSS media for 20 hours. Total cellular peroxide levels were assessed by FITC fluorescence, following incubation with 10 µM CM- H_2_DCF-DA. Unpaired two-tailed Student’s t-test were used to determine p-values. H. LNAI cells were plated in triplicate in 96-well plates and treated with vehicle (DMSO) or riociguat in FBS or CSS media for 20 hours. Cells levels of cellular hydroxyl and peroxynitrite radicals were assessed by FITC fluorescence, following incubation with 10 µM HPF. Unpaired two-tailed Student’s t-test were used to determine p-values.

Riociguat treatment was accompanied by elevation of the pro-apoptotic protein, cl-PARP, and downregulation of the anti-apoptotic proteins, Survivin and Bcl-2 protein, under AD but not androgen-replete conditions (**Fig. 4E**). Thus the loss of viability by riociguat treatment is consistent with induction of AD-induced apoptosis, and consistent with previous reports that elevated cGMP in cancer cells and associated downstream signaling induces apoptosis^33, 34^. We have reported that AD acutely alters cellular reactive oxygen species (ROS) levels^7^ and therefore we assessed whether riociguat treatment affected ROS levels. We found that riociguat minimally affected total cellular superoxide or hydroperoxides levels, as measured respectively by the cell-permeant fluorescent dyes, hydroethidium (HEt) and chloromethyl-dichlorofluorescein diacetate (CM-DCFDA) (**Fig. 4F, G**). However, riociguat treatment increased total levels of high-potency radicals (hydroxyl radical, peroxynitrite), in AD but not in androgen replete conditions, as measured by the cell-permeant fluorophore hydroxyphenyl fluorescein (HPF) (**Fig. 4H**). Acute hydrogen peroxide treatment confirmed that the two dyes measure different species; the DCFDA fluorescence is the same in FBS and CSS as the directly measured species is equivalently added to both (**Fig. 4G**). However, the levels of HPF signal in the hydrogen peroxide-added samples are higher under CSS, consistent with the acutely added hydrogen peroxide feeding into a Fenton reaction and leading to increased hydroxyl radicals (**Fig. 4H**). Superoxide and hydrogen peroxide are largely signaling entities in cancer cells and promote oncogenic signaling^35^ whereas high potency radicals are short-lived, cell-damaging species^36^.

Because high potency radicals can readily react with DNA and produce genotoxic breaks, we next assessed whether increased DNA breaks are associated with riociguat-induced cytoxicity in androgen-deprived LNAI cells. To do so, we used the alkaline single-cell gel electrophoresis (“comet”) assay to assess such damage in cells under sub-cytotoxic treatment conditions. Induction of DNA breaks is visualized as cells with increased DNA “tail” moments under electrophoretic conditions, whereas non-damaged cells visualize as intact spheres (**Supplementary Fig. S4E,** CC3 positive controls vs. CC0 negative controls). However, our results do not implicate DNA breaks as the source of riociguat cytotoxicity via the increased high-potency radicals (**Supplementary Fig. S4F**). Thus, the vulnerability of these cells from AD-induced elevated sGC signaling results from stresses evoked by something other than genotoxic damage. Nevertheless, our results collectively indicate that riociguat treatment elevates damaging ROS species to trigger AD-induced apoptosis in CRPC cells, through as-yet unknown mechanisms.

### The sGC complex Is oxidatively inhibited in CRPC cells and functionally regenerated by AD-induced redox protective responses

Our results in **Fig. 4A, B** suggest that AD enhances sGC signaling. The sGC complex is known to be functionally inactivated, through oxidation of its ferrous heme to a ferric heme^37^, in several inflammatory and age-associated pathologies^20^. However, this phenomenon has not previously been identified as a mechanism of functional sGC modulation in human tumors. We hypothesized that the sGC complex is oxidatively inhibited in CRPC cells relative to CSPC cells, making the former less responsive to riociguat stimulation. Therefore, we tested whether the sGC complex is oxidized in CRPC by using cinaciguat, which stimulates sGC in the absence of a functional heme group^38^. We found LNAI cells showed a robust apoptotic response to cinaciguat treatment even under androgen-replete (FBS) conditions (**Figs. 5A. B, Supplementary Fig. S5A**). A similar response to cinaciguat under androgen-replete conditions was seen in 22Rv1 cells **(Supplementary Fig. S5B**). Given that cinaciguat stimulates only oxidized sGC, it was unable to stimulate cGMP production in CSPC LNCaP SB0 cells (**Fig. 5C**). Thus, CSPC LNCaP cells (unlike LNAI) sustain reductively functional sGC. Significantly, cinaciguat was unable to stimulate cGMP production in the earliest CRPC SB5 variant but was able to progressively stimulate cGMP levels with greater efficacy in the more CRPC-like SB5X and SB5XX variants (**Fig. 5C**). This result indicates that the regulation of sGC signaling progressively shifts from dysregulation of heterodimer formation through decreased β1 subunit in emergent CRPC variants (**Fig. 2**) towards oxidation-induced inactivation of the functional complex in established CRPC cells. We also find that levels of the chaperone protein, HSP90, which binds and stabilizes the apo-β1 subunit to facilitate formation of the stoichiometric complex^39^, are highest in LNAI and lowest in the early emergent variants (**Fig. 5D)**. It is thus possible that the progression from CSPC to CRPC reflects disruption of the maturation process of sGC^40^.

**Figure 5.**
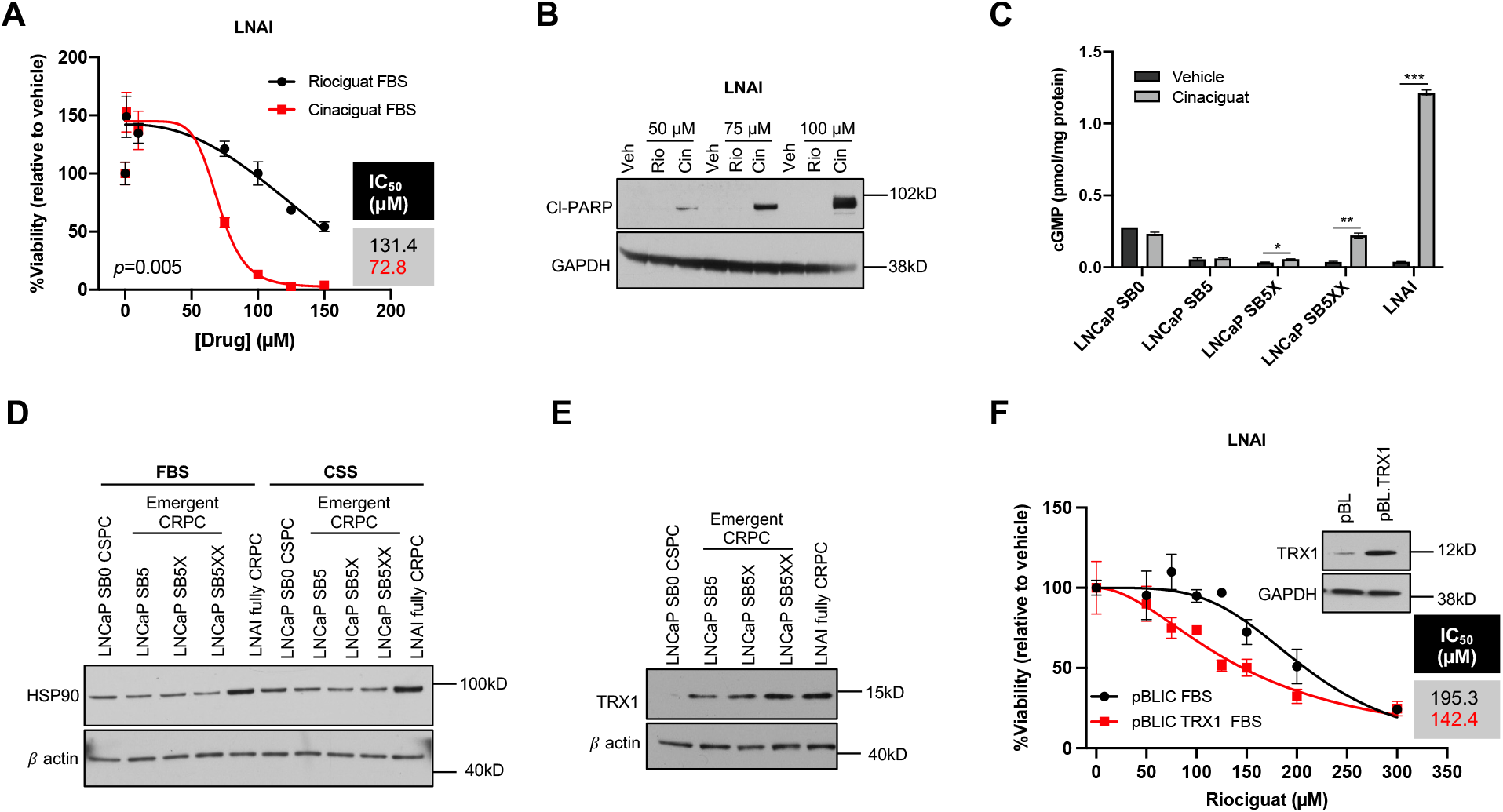
The sGC complex is oxidatively inactivated in CRPC cells and functionally regenerated by AD-induced redox protective response. Note that all error bars represent ± SEM and * p< 0.05, ** p≤ 0.01, *** p≤ 0.001, **** p≤ 0.0001. Note that all immunoblots show the molecular weight of the markers run on the original membrane on the right. A. Cinaciguat decreases the viability of LNAI CRPC cells even under androgen-replete conditions. Cells were plated at 96- well plates in triplicate and treated with riociguat or cinaciguat in FBS media for 72 hours with daily redosing. Cell viabilities, IC_50_s and p-values were established as in Fig. 4C. B. Cinaciguat induces apoptosis in LNAI CRPC cells. LNAI cells were plated in 10 cm dishes under the conditions in (A), and harvested following treatment. Total protein lysates (15 µg) were immunoblotted and probed with the indicated antibodies. C. Cinaciguat stimulates sGC activity in LNAI CRPC to a greater extent than in LNCaP CSPC cells. LNCaP SB0, SB5, SB5X, SB5XX and LNAI cells were plated in FBS-supplemented media, and cGMP levels were measured following incubation with either vehicle or 20 µM cinaciguat for 30 minutes. An unpaired two-tailed Student’s t-test was used to determine p-values. D. Western blotting was carried out using total protein lysates (10 µg) from SB0, SB5, SB5X, SB5XX and LNAI cells cultured under FBS- or CSS-supplemented media for 30h. Blots were probed for HSP90 with β actin as the loading control. E. Levels of the redox-protective thiol protein, TRX1, increase with castration resistance. Western blotting was carried out using total protein lysates (15 µg) from SB0, SB5, SB5X, SB5XX and LNAI cells. Blots were probed for TRX1 with β actin as the loading control. F. TRX1 overexpression sensitizes LNAI CRPC cells to riociguat under androgen-replete conditions. Generation of control vector and TRX1-overexpressing LNAI counterpart lines was validated by immunoblotting (inset). The pBL- and pBL.TRX1-expressing LNAI cells were plated as described in A, and treated with riociguat for 72 hours in FBS-supplemented media, with daily drug redosing. Cell viability and IC_50_s were established as in Fig. 4C.

To further support that sGC is oxidatively inactivated in CRPC cells, we found that the sensitizing effects of AD culture to riociguat treatment could be recapitulated in androgen-replete conditions by culturing LNAI cells at 5% ambient oxygen, which is known to reduce intracellular ROS levels^7, 41–43^ (**Supplementary Fig. S5C**). Thus, sGC is functionally regenerated under conditions that decrease cellular oxidative stress. We previously reported that CRPC cells elevate the redox-protective protein, TRX1, to protect against AD-induced ROS levels and cell death^7^. TRX1 is a known mechanism for restoring sGC activity via thiol reduction ^44–46^. We found TRX1 levels progressively increased when comparing its levels across the continuum of progressively CRPC matched cell lines (CSPC LNCaP to emergent CRPC SB5, SB5X, SB5XX to fully CRPC LNAI) (**Fig. 5E**). When we overexpressed TRX1 in LNAI cells (**Fig. 5F**), we found the TRX1- overexpressing counterparts exhibited a better riociguat response relative to control cells, even under androgen-replete conditions (**Fig. 5F**). The degree of sensitization in FBS was not equivalent to the sensitization produced by AD (**Fig. 4C**), suggesting there are likely other redox-protective mechanisms evoked by AD in CRPC cells that reductively regenerate the sGC complex.

Regardless, these results collectively point to oxidative inactivation of sGC as a critical regulatory mechanism in advanced CRPC, and that the redox-protective responses generated by CRPC to protect against AD-induced death create a paradoxical therapeutic vulnerability to riociguat. Collectively, these data support that the sGC complex is abrogated via decreased β1 expression levels (**Fig. 2A, B**) in emergent CRPC cells whereas progression to full-blown CRPC is associated with restoration of the sGC heterodimer but inhibition of its activity through heme oxidation and potentially thiol oxidation.

### Riociguat treatment decreases CR tumor growth through on-target stimulation of sGC activity and AD-induced apoptosis

Because AD conditions sensitized LNAI cells to riociguat, we next tested whether riociguat treatment combined with in vivo AD would restrain CRPC growth. To this end, we subcutaneously injected 2 million LNAI cells into the flanks of 6 week old Nu/Nu castrated male mice, which lack adrenal androgens^47^ and thus are systemically depleted of androgens. Following palpable tumor growth, animals were randomized into groups that received intraperitoneal (IP) injections of either vehicle (DMSO) or 20 mg/kg/day riociguat daily. This dose was well tolerated as riociguat-treated animals did not exhibit weight loss (**Supplementary Fig. S6A)** or other physical signs of drug toxicity. Riociguat-treated animals showed both reduced tumor growth kinetics as well as endpoint tumor sizes and tumor weights (**Figs. 6A, B; Supplementary Fig. S6B, C**). Hematoxylin & eosin staining (H&E) indicated a perceptible reduction of the pan-proliferative marker, Ki67, in riociguat-treated tumors relative to vehicle-treated (**Fig. 6C**) as well as morphology consistent with cell death (reduced cellularity, pyknosis; **Fig. 6C**). Indeed, riociguat-treated tumors presented with a striking increase in apoptosis-associated DNA fragmentation, measured via TUNEL staining in formalin-fixed tumor sections (**Fig. 6D**). These tumors also had higher protein levels of the apoptotic marker, cleaved PARP (cl-PARP) relative to control lysates (**Fig. 6E**). Riociguat treatment did not impair the ability of PSA levels to correlate positively with castration-resistant tumor growth (**Supplementary Fig. S6D**).

**Figure 6.**
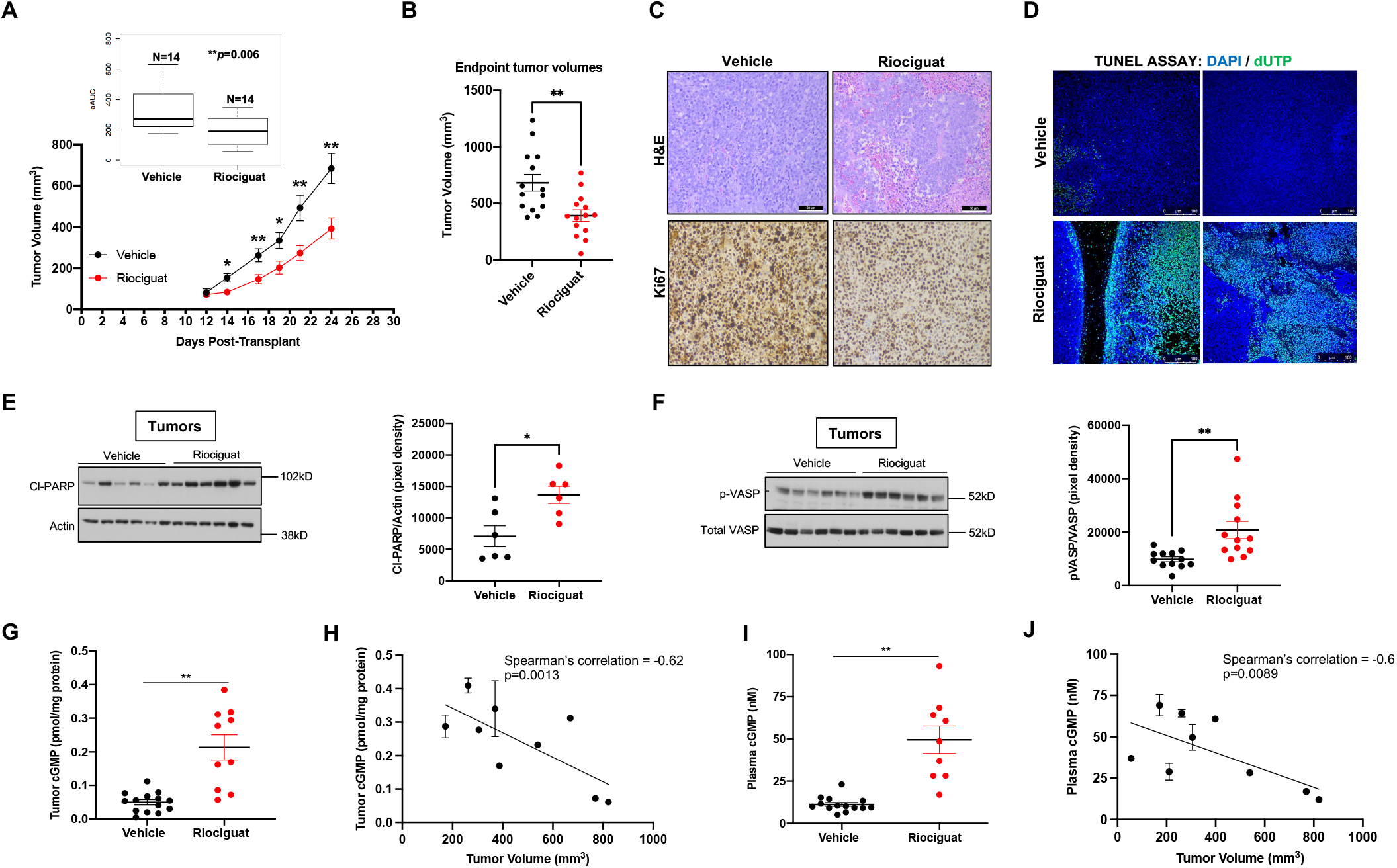
Riociguat treatment decreases CR tumor growth through on-target stimulation of sGC and resensitization to AD-induced apoptosis. Note that all error bars represent ± SEM and * p< 0.05, ** p≤ 0.01, *** p≤ 0.001, **** p≤ 0.0001. Note that all immunoblots show the molecular weight of the markers run on the original membrane on the right. A. Riociguat reduces castration-resistant tumor growth. LNAI tumor-bearing castrated male Nu/Nu mice were randomized into groups receiving either daily IP injections of riociguat (20 mg/kg/day) or DMSO. Tumor growth was monitored three times weekly. Mean tumor volumes per treatment group across the 24-day experiment are shown. Adjusted areas under the curve (aAUCs) are plotted within the inset for the vehicle group (n=14) and the riociguat-treated group (n=14). Wilcoxon two-sample test was used to determine p-values. B. Endpoint tumor volumes (mm^3^) in each treatment group (n=14) are shown. Each point represents an individual tumor/mouse. Unpaired two-tailed Student’s t-test was used to determine p-values. C. Representative H&E and Ki67-stained immunohistochemical images for tumors from the vehicle and riociguat-treated groups. The size bar, in white, represents a 50 μm scale. D. TUNEL staining shows riociguat treatment induces apoptosis. The TUNEL assay was carried out on tissue sections from vehicle and riociguat-treated tumors. Representative images of co-localized dUTP/DAPI staining are shown for two separate representative tumors in each indicated group. Images were acquired through identical exposures per channel. The size bar represents a 100 μm scale. E. Western blotting for the apoptotic marker, cleaved-PARP (cl-PARP), in xenograft tumor lysates (15 µg) from vehicle or riociguat-treated mice (left). Actin is used as the loading control. Quantitation of the cl-PARP immunoblot signal from the respective indicated tumor lysates, normalized to actin, is shown (Right). Unpaired two-tailed t-test with Welch’s correction was used to determine p- values. F. Immunoblotting for phospho (Ser239)-VASP in tumor lysates (30 µg) from vehicle or riociguat-treated mice. Total VASP levels are shown as loading control. Quantitation of the p-VASP protein signal, normalized to total VASP, is shown (right). Unpaired two-tailed t-test with Welch’s correction was used to determine p-values. G. Intratumoral cGMP levels were measured in tumor lysates from the vehicle or riociguat tumor groups. Unpaired two-tailed t-test with Welch’s correction was used. H. Intratumoral cGMP levels from riociguat-treated animals correlate inversely with the tumor volumes. Spearman correlation test values are shown. I. Plasma cGMP levels were measured in the vehicle or riociguat groups. Unpaired two-tailed t-test with Welch’s correction was used to determine p-values. J. Plasma cGMP levels from riociguat-treated animals correlate inversely with xenograft tumor volumes. Spearman correlation test values are shown.

To assess whether riociguat was acting through its anticipated mechanism and stimulating sGC catalytic activity, we assessed levels of phosphorylated vasodilator-stimulated protein (p- VASP), a downstream marker for cGMP-dependent signaling via protein kinase G^48^. As anticipated, the treated tumors had significantly higher p-VASP protein levels (**Fig. 6F**). We next measured cGMP levels in both lungs (the site of maximal bioactivity) and in the tumors. While cGMP levels were significantly elevated in both the lungs as well as in the tumors of the treated animals (**Supplementary Fig. S6E**, **Fig. 6G**), the cGMP levels and their relative variability among intratumoral control tumors was far less than seen in the corresponding lung tissue from control animals. This is consistent with human xenograft CRPC tumors possessing dampened baseline (unstimulated) sGC activity compared to normal murine lung tissue.

We further found that cGMP levels showed a significant inverse correlation with castration-resistant growth (**Fig. 6H**), supporting both on-target anti-tumor activity of riociguat as well as prognostic potential for cGMP levels in addition to PSA. We also measured cGMP levels in plasma, which can be obtained noninvasively compared to tumor tissue, and found these levels were higher in the riociguat-treated animals and also correlated negatively with tumor growth (**Fig. 6I, J**). Indeed, plasma cGMP also showed a positive correlation with intratumoral cGMP levels (**Supplementary Fig. S6F**) and a negative correlation with PSA (**Supplementary Fig. S6G**). Thus measuring plasma cGMP levels, a non-invasive approach, is a similarly effective biomarker for riociguat treatment response and castration-resistant tumor growth as the intratumoral levels. Collectively, our data herein show that riociguat effectively suppresses castration-resistant tumor growth by inducing apoptosis, and that the degree of its on-target efficacy can be readily measured by declining PSA as well as by increasing intratumoral or plasma cGMP levels.

### Tumor-inhibitory responses to riociguat are associated with improved oxygenation and enhanced sensitivity to radiation

Castration induces hypoxia^49^ and hypoxic signaling promotes prostate cancer progression/metastasis and aggressive phenotypes^50, 51^. Targeting hypoxic cells inhibits PC progression^52^. Because sGC stimulation is expected to increase tissue oxygenation as per its physiologic function, we determined whether its PC-inhibitory function arose from its abrogation of hypoxia. To assess if riociguat treatment alters CRPC oxygenation, two hours prior to euthanasia, we randomly assigned experimental animals (vehicle and riociguat) for injection with the hypoxia sensor, pimonidazole (Hypoxyprobe, 60 mg/kg), which accumulates in hypoxic tissues. Hypoxyprobe is detectable in formalin-fixed sections via a fluorescent secondary antibody^53, 54^. We found riociguat-treated sections showed a striking decrease in Hypoxyprobe signal vs. their vehicle counterparts, regardless of tumor size (**Fig. 7A**, images representative of n=5 animals/group). This decrease in hypoxia was irrespective of tumor size as all riociguat-treated tumors assessed (across a range of tumor volumes) showed lower staining relative to their size-matched vehicle-treated counterparts (**Fig. 7A**, representative images). Immunohistological staining for the endothelial cell marker, CD31, showed poorly formed, constricted vessels in the vehicle-treated tumors. By contrast, the riociguat-treated tumors all showed organized staining around an open lumen in the riociguat-treated tumors (**Fig. 7B**), indicative of improved vasculature. Given that CRPC are known to have leaky and dysfunctional vasculature as well as vasoconstriction due to aberrant NOS activity^55, 56^, the tumor oxygenation observed with riociguat treatment likely arises from this vascular reorganization or canalisation of tumor blood vessels.

**Figure 7.**
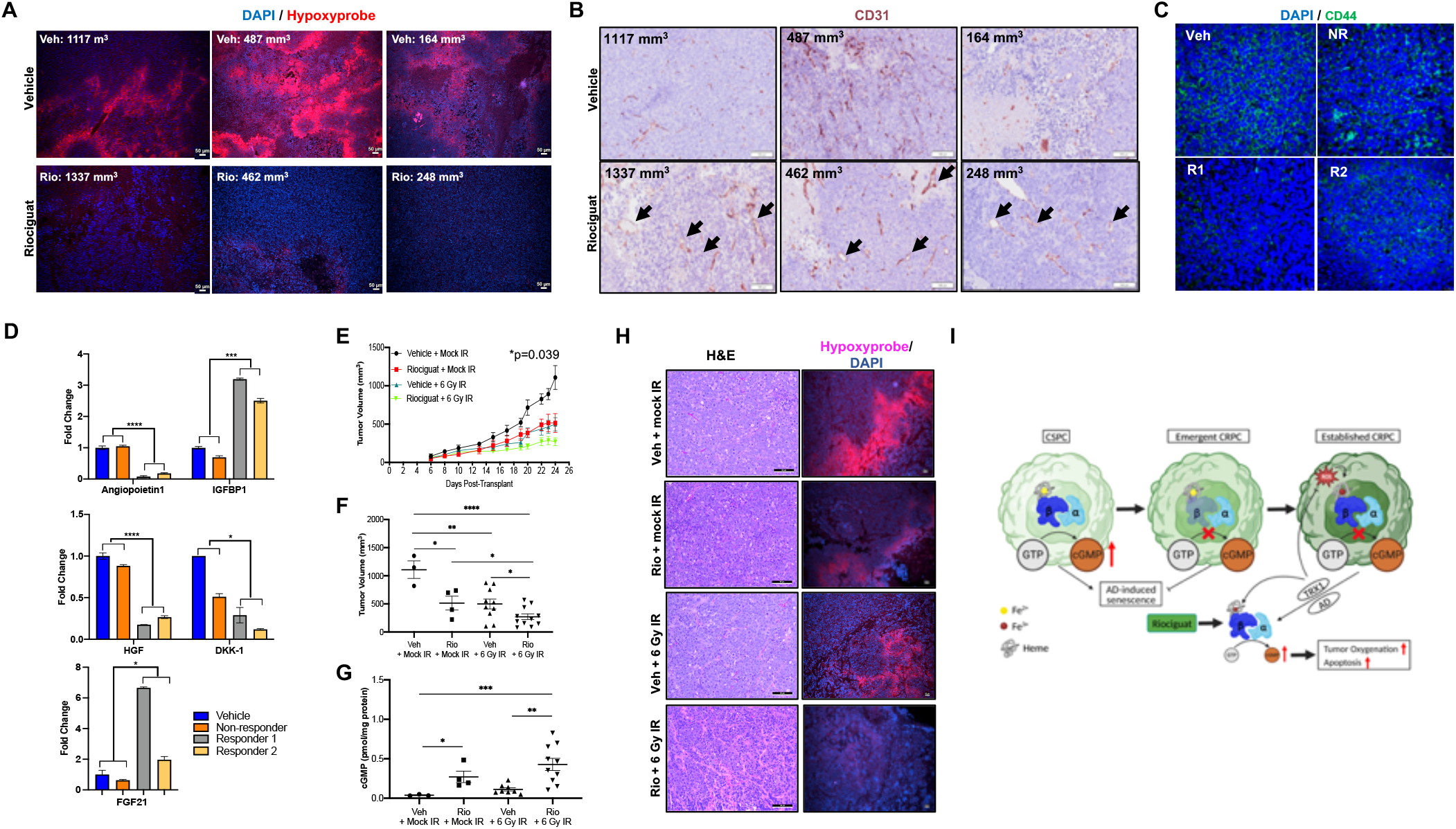
Tumor-inhibitory responses to riociguat are associated with improved oxygenation and enhanced sensitivity to radiation. Note that all error bars represent ± SEM and * p< 0.05, ** p≤ 0.01, *** p≤ 0.001, **** p≤ 0.0001. A. Riociguat treatment oxygenates CRPC tumors. Representative immunofluorescent images of vehicle- or riociguat-treated tumors stained with pimonidazole (Hypoxyprobe) and counterstained with the nuclear dye, DAPI. Endpoint tumor volumes are indicated within each panel. The size bar represents a 50 μm scale. B. Riociguat treatment induces tumor vascular reorganization. Representative immunohistological images of the same tumors from (A), stained for the vascular endothelial marker, CD31. Arrows indicate canalised blood vessels. The size bar represents a 100 μm scale. C. Anti-tumor response to riociguat treatment correlates with loss of the PC stem cell marker, CD44. Representative immunofluorescent images of a vehicle-treated, a riociguat-treated nonresponsive tumor (NR), and two riociguat-responsive (R1, R2) tumors stained for CD44, shown in green, and counterstained with DAPI (blue). D. Cytokine markers of anti-tumor response to riociguat treatment. Sera obtained from vehicle, non-responder, responder 1, and responder 2 animals in (C) were assayed for systemic changes in host (murine) cytokine levels. The indicated cytokines fit the criteria of being the only ones where the non-responder and vehicle-treated values are similar and where the responder 1, responder 2 values trend in the same direction. Each cytokine value is the average of n=2. Results are represented as fold-change relative to the values established from vehicle-treated sera. The p values compare changes in group 1 (vehicle, non-responder) vs those in group 2(responder 1, responder 2), and were established using unpaired two-tailed Student’s tests. E. Riociguat sensitizes CRPC tumors to radiation. LNAI tumor-bearing castrated male Nu/Nu mice were randomized into groups receiving daily IP injections of riociguat (20 mg/kg/day) or vehicle (DMSO) for 96 hours. Following this, animals were further randomized in each group to receive either a single dose of 6 Gy ionizing radiation (IR) or mock irradiation (0 Gy). Daily IP injections of vehicle or riociguat were continued for the duration of the experiment. Mean tumor volumes per treatment group across the 24- day experiment are shown: vehicle/mock IR (n=3), riociguat/mock IR (n=4), vehicle/6Gy IR (n=10), riociguat/6Gy IR (n=11). The p values shown are comparing aAUC of the curves (vehicle/6 Gy IR vs riociguat/6 Gy IR) using a Kruskal-Wallis test. F. Endpoint tumor volumes (mm^3^) from all treatment groups. Each point represents an individual tumor. Unpaired two-tailed Student’s t tests were used to determine p-values. G. Intratumoral cGMP levels were measured from xenograft tumors across all treatment groups. Unpaired two-tailed t-tests with Welch’s correction were used to determine p-values. H. H&E immunohistology is shown alongside co-localized Hypoxyprobe/DAPI immunofluorescent staining from representative tumors across all treatment groups. I. Overview of the role for sGC signaling in prostate cancer progression, showing the loss of sGC activity in the CSPC-to-CRPC transition through dysregulated heterodimer formation to evade ADIS, and through oxidative inactivation in established CRPC.

Because hypoxic niches house tumor stem cells known to promote CRPC^57, 58^, we also stained for CD44, a key PC stem cell marker^59, 60^. We found that lower CD44 staining correlated with riociguat response, with greater decrease in staining seen in smaller riociguat-treated tumors (responders – R1, R2, less than 500 mm) (**Fig. 7C**). By contrast, a non-responder treated tumor (NR, similar in size to vehicle-treated) did not show similarly decreased CD44 (**Fig. 7C**). When we evaluated changes in the host paracrine environment via a murine cytokine panel, very few cytokines were changed oppositely in the riociguat responders vs. the nonresponder and vehicle-treated animals. Of these significant changes, two were associated with improved vasculature and reduced hypertension (**Fig. 7D**, topmost graph): Angiopoietin-1, synthesized by vascular smooth muscle cells, is a neovascularization factor, whose high plasma levels are associated with pulmonary hypertension^61^. Its decrease in the riociguat responders is consistent with the lack of observed neo-angiogenesis (**Fig. 7B**). Similarly, low circulating levels of IGFBP1 are reported to lower blood pressure and increase nitric oxide production^62^, suggesting this is another marker of riociguat efficacy. The other three host cytokines, altered differentially in responders vs. nonresponder/vehicle, were associated with decreased stemness **(Fig. 7D**, middle and bottom). These included elevated FGF21, which improves vascular function^63^ and promotes apoptosis in prostate cancer cells^64^, and decreased HGF and DKK1, both factors associated with anti-differentiation functions and tissue regeneration^65, 66^ (**Fig. 7D**).

Thus, our results show that, in addition to increasing CRPC apoptosis and loss of tumor self-renewal markers, riociguat treatment modulates the tumor microenvironment, including tumor oxygenation, vascular reorganization and altered paracrine factors implicated in revascularization and inhibition of self-renewal. Despite variability in its CRPC-inhibitory effect, riociguat oxygenates even non-responder tumors (**Fig. 7A**), creating a vulnerability to additional treatment modalities. Castration-resistant tumors become resistant to radiation therapy^67^ in large part due their hypoxic state^68, 69^. It is reported this resistance to radiotherapy is mediated at least in part by the preservation of self-renewing prostate cancer stem cells in hypoxic niches^70–72^. Therefore, due to its destruction of hypoxic tumor niches, we tested whether riociguat treatment would increase the efficacy of ionizing radiation (IR) in xenograft CRPC tumors. Combining riociguat with IR significantly improved on tumor growth suppression over radiation or riociguat alone, with combined radiation and riociguat showing the greatest suppression of tumor growth relative to untreated (**Figs. 7E, F, Supplementary Fig. SS7**). We also noted that riociguat-mediated tumor growth inhibition was comparable to radiation alone (**Figs. 7E, F**). CRPC inhibition under these conditions was paralleled by the highest induction of sGC activity, as evaluated by cGMP levels, in combined riociguat/IR-treated tumors (**Fig. 7G**). We hypothesized that IR sensitization would occur via riociguat’s ability to reduce tumor hypoxia. Therefore, we again evaluated intratumoral hypoxia using Hypoxyprobe. We found tumor oxygenation was even more marked in the combination riociguat/IR cohort compared to riociguat alone, which coincided with reduced tumor cellularity as visualized through H&E (**Fig. 7H**). Therefore, our results support combining riociguat with IR to enhance the efficacy of radiotherapy in CRPC.

## Discussion

Here we describe the first studies rationalizing a therapeutically actionable role for sGC agonists, notably the FDA-approved vasodilator riociguat, in reducing CRPC burden and in enhancing efficacy of radiation therapy through tumor oxygenation. Our results support that sGC signaling is dysregulated in the progression from CSPC to CRPC by disrupted stoichiometry of the sGC heterodimeric complex and inhibited in established CRPC by sGC oxidation.

We provide that first evidence that the sGC complex activity enforces the AD-induced senescence arrest in CSPC, required to restrain progression to CRPC. Indeed, sGC activity is nearly undetectable in emergent CRPC cells under AD conditions due to loss of expression of the β1 subunit, which is essential for sGC complex activity. Mimicking this effect through β1 CRISPR knockout leads to a decreased number of AD-induced senescent CSPC cells and facilitates castration-resistant tumor growth. Thus, our studies support that the sGC complex needs to be fully suppressed to effectively lower the ADIS barrier to CRPC emergence.

Significantly, acquisition of the established CRPC state is associated with restoration of β1 expression and sGC heterodimer formation. However, at this stage, sGC activity (and cGMP production) is downregulated through oxidative inactivation of the sGC complex, underscoring the overall tumor-suppressive role of sGC signaling activity in CRPC. This functional inactivation does not appear to be a binary process but rather there seems to be a population-based shift from cells, which have suppressed sGC heterodimer formation through decreased β1 subunit expression, to cells which have sufficiently restored β1 expression to form the stoichiometric complex but with an oxidatively inactivated heme. Thus, the SB5 or SB5X early CRPC cells do not show any stimulation by either riociguat or cinaciguat (indicating loss of the heterodimer), but the more CRPC-like SB5XX cells (which do not respond to riociguat) **(Fig. 2C, Fig. 4A, B)** do indeed show a detectable increase in cinaciguat-induced cGMP, suggestive of oxidative sGC inactivation **(Fig. 5C**).

The combination of sGC transcriptional dysregulation in emergent CRPC cells not typically found in patient specimens, and the protein oxidation-induced functional inhibition in established CRPC cells means that the changes we identified here cannot be easily captured in patient transcriptomics datasets. However, these changes are reflected in decreased sGC activity as measured by cGMP levels, either intracellularly, intratumorally or in sera/plasma samples. Indeed, cGMP levels are lower in both preclinical and clinical CRPC vs. CSPC samples. Strikingly, evaluation of cGMP levels in the sera of patients who progressed to CRPC suggests decreased cGMP in CRPC vs. counterpart CSPC specimens has the potential to predict aggressive disease and mortality (**Fig. 1J**).

We previously reported that CRPC cells mount a redox-protective response, notably elevated TRX-1, to protect against AD-induced death^7^. Here we showed that this response paradoxically sensitizes CRPC cells to riociguat, ostensibly by reductively regenerating the sGC complex. This is seen by the increased sensitization of LNAI cells to riociguat under AD culture and treatment with the gold-standard anti-androgen enzalutamide (**Fig. 4C, D**). Indeed, TRX1- overexpressing cells are more sensitized to riociguat-induced cytotoxicity compared to control cells, even under androgen-replete conditions (**Fig. 5F**). This is consistent with other studies reporting TRX1 as a critical factor for keeping sGC in its reduced functional state^44, 45^. This redox Achilles Heel of CRPC cells can likely be exploited by other drug mechanisms. However, we note that agents that sensitize CRPC to AD such as the TRX1 inhibitor, PX-12, or the TRX-R inhibitor, auranofin, will likely antagonize the tumor-inhibitory effects of riociguat by removing AD redox-protective effects. However, other reductive agents with clinical potential, such hydrogen sulfide (H_2_S)^73^, known to reduce sGC^74^, may be able to enhance the action of riociguat in CRPC.

The fact that the sGC subunit transcription is maintained in established CRPC and the loci are not deleted suggests that sGC function may be focally required at a low baseline level for physiologic purposes, possibly related to vascular structure and function. In breast cancer, suppression of the β subunit has been shown to occur through epigenetic means and treatment with both selective and non-selective HDAC inhibitors can restore expression of sGCβ1^11^. Further studies will be required to ascertain why and how the sGC complex is both transcriptionally and oxidatively dampened or its maturation impaired in prostate cancer. It has been reported that FOXO family proteins, whose dysregulation occurs in PC^75, 76^, transcriptionally control sGC expression^77^ and particularly expression of the β subunit^78^. Moreover, we have shown that inappropriate AR activation in the low ligand environment of CRPC produces ROS^7^, which can reversibly inactivate sGC^37^. Thus, sGC activity appears to be a rheostat for progression to CRPC rather than an absolute barrier. This allows the sGC complex to be an effective therapeutic target in CRPC, as riociguat can stimulate cGMP production even from low levels of the sGC complex.

High levels of cGMP have previously been reported to kill cancer cells. Activation of PKG via cGMP through the sGC activator YC-1 and addition of the phosphodiesterase-resistant cGMP homolog, 8-bromo-cGMP, to MCF-7 and MDA-MB-468 breast cancer cells induces apoptosis through caspase activation^34^. Elevation of cGMP in head and neck squamous cell carcinoma (HNSCC) through NO donors, sGC stimulators and PDE5A inhibitors also leads to apoptosis through PKG signaling^79^. Activation of PKG has also been reported to be sufficient to induce apoptosis in SW480 colorectal cancer cells^33^. We find that riociguat-induced cytotoxicity is associated with lower levels of the caspase-inhibitory proteins, Bcl2 and the IAP, Survivin (**Fig. 4E**). We also show that the in vivo anti-tumor activity of riociguat in CRPC is associated with elevated levels of p-VASP, the downstream target of PKG (**Fig. 6F**). Given the oxidative regulation of sGC activity, we anticipated riociguat-induced stimulation would involve ROS-dependent effects. While stimulation of riociguat in CRPC cells does not significantly alter superoxide or hydrogen peroxide levels, it does increase high potency radicals (peroxynitrite, hydroxyl radical) under AD (**Fig. 4F-H**), which are short lived and highly damaging. Indeed, the changes we evaluated using fluorophores in our cells are likely to be underestimates of the true extent to which sGC stimulation under AD generates these highly damaging radicals. Both the source(s) of these ROS and the actual mechanism of cell death still remain to be established. More broadly, the observed increase in HPF fluorescence suggests that AD may also increase labile metals such as iron and copper, which catalyze production of these high potency radicals.

The sGC-cGMP pathway has been previously evaluated as a target in prostate cancer^80–82^. These studies propose inhibition rather than stimulation of sGC signaling as a tumor-repressive strategy. However in vivo experiments with sGC-inhibited CRPC lines were carried out in non-castrated animals, thus losing the critical clinically relevant effects of systemic AD, or were carried out using a CSPC line, VCaP^82^, which does not appear to show any dependency on the sGC subunits (**Supplementary Fig. S1B,C**). Furthermore, sGC was inhibited using compounds based on the oxidizing agent 1-H-[1,2,4]oxadiazolo[4,3-a]quinoxalin-l-one (ODQ), which is known to induce cytotoxicity via sGC-independent nonselective effects on multiple heme proteins^83, 84^. Our studies robustly support that riociguat functions through on-target mechanisms in reducing CRPC growth, supported by the β1 subunit KO experiments and the inverse correlation between cGMP levels and tumor sizes.

Prior studies indicate high levels of NO (the sGC ligand) have anti-CRPC effects. NO donors or high endothelial nitric oxide synthase (eNOS) expression inactivate AR via s-nitrosation^85^, and reduce tumorigenesis by CRPC cells^85, 86^. Indeed, several limitations noted for NO donors, including unpredictable rate of release, lack of selectivity and induced vascular resistance, do not pertain to sGC agonists^87^. Furthermore, unlike NO donors, sGC agonists are not a source of systemic nitrosative stress, which can induce tissue damage and contribute to oxidative inactivation of the sGC complex, preventing on-target stimulation. Furthermore, adverse systemic effects of NO are thought to be largely cGMP-independent^88^. NO donors likely work by nitrosative inactivation of AR^85^, a vulnerability in advanced PC which then adaptively suppresses NOS activity limiting physiologic sGC stimulation. Further, the observed oxidation of sGC in CRPC limits the on-target utility of NO donors due to inactivation of the catalytic heme. Riociguat, however, acts on sGC independently of NO levels, but can also sensitize sGC to low NO concentrations^26^ to elevate cGMP levels. Use of PDE5 inhibitors^89^ such as sildenafil and tadalafil can also elevate cGMP by preventing its hydrolysis, and these inhibitors are also clinically actionable. However, sGC is likely to be the main determinant of intratumoral cGMP levels in CRPC as PDE5A levels appear to be low in PC^19^ (and **Supplementary Fig. S1G**). Therefore, whether PDE5 inhibitors can improve or enhance the effects of sGC agonists, and whether they possess therapeutic benefit in some prostate cancer settings remains to be investigated.

CRPC tumors are known for leaky and disorganized vasculature and a hypoxic tumor microenvironment^49, 55, 56^. Hypoxic tumor niches potentially house CRPC-driving PC stem cells^90^. These cells reportedly drive CRPC emergence and maintenance and are marked by high CD44 expression^91, 92^. Thus, the elimination of high CD44 cells by riociguat has the potential of increasing AD efficacy and indeed shortening its duration. Additionally, the ability of riociguat to canalise existing vasculature has additional therapeutic advantages via improved tumor oxygenation. Indeed, radiation efficacy, which is compromised by hypoxia in the advancing PC tumor, is restored by riociguat induced-tumor oxygenation. CSPC tumors are very sensitive to radiotherapy, making it an effective standard-of-care approach at this disease stage^93^. Riociguat treatment appears to restore this aspect to CRPC tumors in addition to resensitizing them to AD. Riociguat induces oxygenation in large as well as small tumors despite highest cGMP levels correlating significantly with smaller tumors (**Fig. 6H,J, Fig. 7A**). One explanation of this observation is that riociguat bioavailability through our IP procedure is sufficient for raising cGMP levels to promote oxygenation (a known acute effect of sGC signaling) but not apoptosis, which may occur only among the tumors with the highest cGMP increases upon riociguat treatment (**Fig. 6G, I**). Nevertheless, the consistent tumor oxygenation by riociguat opens the way for improving efficacy of radiotherapy and possibly other drug regimens, due to the improved canalisation of vasculature.

Although the observed anti-tumor response via sGC stimulation has clear cell-intrinsic components, we also noted that the host cytokine profiles in riociguat-treated animals with greater reduction in tumor growth (responders) vs. those with lesser reduction (non-responders) show several differences (**Fig. 7D**). Indeed, despite undergoing riociguat treatment, the animals with non-responsive tumors have a cytokine profile very similar to the vehicle-treated animals. As riociguat-induced changes support vascular remodeling and loss of cellular plasticity, it is possible these paracrine responses are consequences rather than causes of riociguat-induced anti-tumor responses. Nevertheless, these changes have the potential to serve as non-invasive markers of riociguat treatment efficacy in future patient-centered trials.

Collectively, our study points to sGC signaling as a key barrier to castration resistance through its reinforcement of AD-induced senescence (**Fig. 7I).** This is, to the best of our knowledge, the first report of sGC signaling in reinforcing tumor-suppressive senescent phenomena. The stoichiometric restoration of sGC heterodimer formation in fully established CRPC and its reductive regeneration under androgen deprivation provides a therapeutic opportunity to stimulate sGC-induced apoptosis by repurposing riociguat. While riociguat treatment is able to increase cGMP levels to a greater extent in some tumors than others in our study, the uniform tumor oxygenation through vascular canalisation of treated tumors opens the way for improving other standard-of-care treatments, for instance radiation. In patients, a decline in sGC signaling, evaluated through systemic cGMP levels, portends worse disease prognosis and mortality following progression to CRPC. Thus, we describe here a new biological pathway involved in the etiology and progression of CRPC as well as a novel actionable treatment for this deadly disease through repurposing riociguat, a safe and well-tolerated vasodilator.

## Materials and Methods

### Patient-derived serum

De-identified matched sera from CSPC/CRPC patients were received from the NCI biospecimen repository (NCT00034216).

### Cell lines

LNCaP (CRL-1740, denoted as LNCaP SB0 in this study) and 22Rv1 (CRL-2505) were obtained from ATCC. LAPC4 cells were a kind gift of Dr. John Isaacs, Johns Hopkins University (Baltimore, MD). None of the cells used in this study are listed in the ICLAC database of commonly misidentified lines. All cell lines used in this study, including our in-house LNCaP CRPC derivatives, LNCaP SB5 variants and LNAI, were validated by short term-repeat (STR) profiling (Genetica Corp). Additionally, all cells used in xenograft animal experiments were certified pathogen-free prior to use. LNCaP cells engineered for disruption of functional sGC activity (sGCβ1 knockout pools) were generated by Synthego Corporation. The sgRNA for sGCβ1(ENST00000264424; sequence: UGCCACACUGAGUGACCACU) targets exon 6. All cell culture reagents described in this section were obtained from Gibco, Life Technologies, except for sera, FBS and charcoal stripped serum (CSS), were obtained from Hyclone. Cell lines were cultured in RPMI-1640 medium (Gibco, 11875093). All media were supplemented with either 5% (LNCaP, LNCaP SB5, LNAI, LAPC4) or 10% (22Rv1) FBS (Hyclone, SH30396) for cell line growth and maintenance. To produce androgen-deprived (AD) conditions, cells were cultured in their appropriate base media supplemented with 5% or 10% CSS (Hyclone, SH30068). To deplete androgen, cultures were washed three times with CSS-supplemented media with 5-minute incubations between washes at 37°C in a humidified incubator. All media were supplemented with 100 U ml^−1^ penicillin/streptomycin (Gibco, 15140122). Cells were cultured at 37 °C in a humidified incubator at 21% oxygen/5% CO_2_ or 5% O_2_/5% CO_2_ (HeraCell Tri-Gas, ThermoFisher Inc).

### Drug treatments and dose response assays

For all cell lines, 750 cells per well were plated in triplicate in 96-well plates in FBS-supplemented culture medium. Twenty four hours after plating, cells were treated with either riociguat (Cayman Chemicals, 9000554) or cinaciguat (Cayman Chemicals, 17468) at various concentrations. For androgen blockade, LNAI cells were pre-treated for 24 hours in 10 μM enzalutamide (Selleckchem, S1250), followed by treatment with various concentrations of riociguat combined with 10 μM enzalutamide or riociguat alone. Drug and media were replenished daily for 72 hours, after which viability was measured using the luminescence-based the CellTiter-Glo® Assay (Promega, G7571) on a SpectraMax iD3 or iDx Microplate Reader (Molecular Devices, LLC). Data were normalized to luminescence values from vehicle-treated controls within each group and plotted as %viability (relative to vehicle). Curve fits were modeled using four-parameter logistic (4PL) regression to derive best fit value, IC_50_. Individual areas under the dose-response curves (AUCs) were calculated and compared by group. All tests comparing aAUC utilized the Wilcoxon test for two groups or the Kruskal-Wallis test for more than two groups.

### DNA constructs and viral transduction

The TRX1 overexpression construct was generated as previously described^7^. Retroviral supernatant production was carried out in HEK 293 T cells (ATCC), and infection of target cells was performed as described previously^7^. Transduced cells were selected in 250 µg ml^-1^ G418 (Life Technologies, 11811031)-containing media, corresponding at a minimum to the time taken for untransduced cells to die completely in selection media. Protein knockdown or overexpression was verified via western blotting.

### Gene expression analysis

Differences in gene expression levels between parental LNCaP SB0 and their CRPC derivative LNCaP SB5, as well as expression of LNCaP SB0 under baseline FBS (proliferating) or CSS (senescent) conditions, were determined through gene expression microarray profiling via the Illumina platform (version HT12). Cells were cultured in either FBS or CSS- supplemented media for 8 days. Equivalent cell numbers (∼3 × 10^6^) across all samples were harvested in QIAzol lysis reagent (Qiagen). Illumina gene expression raw data were transformed using variance-stabilizing transformation (VST) and log2 transformation, and then normalized through quantile normalization using Bioconductor package lumi (v2.24.0). After pre-processing, the limma (v3.28.17) Bioconductor package, which was implemented with a moderated t-test, was used to detect differentially expressed genes between each comparison. The raw p-values of the differential tests were adjusted for multiple testing with Benjamini and Hochberg false discovery rate (FDR) correction. Gene Set Enrichment Analysis (GSEA)^27^ was performed using the GSEA software on pre-ranked gene expression lists. Raw expression values from TCGA and SU2C datasets were obtained from the Human Protein Atlas (https://www.proteinatlas.org/) and cBioportal (http://www.cbioportal.org/) *GUCY1A1*, *GUCY1B1* expression levels were analyzed in GraphPad Prism (v. 9).

Single cell RNA sequencing (sc RNA-seq) data were retrieved and analyzed through single cell portal (https://singlecell.broadinstitute.org/single_cell). The distribution of GUCY1A1 and GUCY1B1 in human prostate tissue were visualized in two dimensions through Uniform Manifold Approximation and Projection (UMAP) plots. The expressions of *GUCY1A1* and *GUCY1B1* were visualized in in dot plot format as generated on the website.

### Quantitative PCR (qPCR) analyses

The mRNA from cultured cells was extracted using the RNeasy (Qiagen, 74136) or RNAqueous-4PCR kit (Life Technologies, AM1914). Complementary DNA was synthesized using 0.5 μg of RNA using the High Capacity cDNA Reverse Transcription kit (Life Technologies, 4368814). The qPCR reaction was set up with 1 μl of diluted complementary DNA and 20X TaqMan probes and TaqMan Universal PCR Master Mix (Life Technologies, 4324018) in a 15 μl total volume. Samples were run in triplicate on an Applied Biosystem (Life Technologies) Real-Time machine using a StepOne program at 95 °C for 10 min and 40 cycles of the following: 95 °C for 15 s and 60 °C for 1 min. Gene expression levels were calculated using the 2^−ΔΔCt^ method. The following gene-specific TaqMan primer/probe sets were used: *GUCY1A1* (Hs01015574_m1), *GUCY1B1* (Hs00168336_m1), *ACTB* (internal normalization control; Hs99999903_m1) (ThermoFisher Scientific).

### Western blotting

Cells in culture were harvested by mechanical scraping on ice and were lysed in a sodium fluoride (NaF) buffer as previously described. Protein lysates from tumors were made using RIPA buffer (ThermoFisher Scientific, 89900), supplemented with a protease inhibitor cocktail (Roche, 11697498001). Protein concentrations were measured using the Pierce BCA Protein Assay kit (ThermoFisher Scientific, 23225). Approximately 10–30 µg of total protein was run on a 4–12% Bis-Tris pre-cast NuPage gel (Life Technologies, NP0321 or NP0322) on the Novex gel system and subsequently transferred onto PVDF membrane (Immobilon, EMD Millipore, IPVH000010) at 35 V at 4 °C. Blots were probed with antibodies against the following proteins: sGCα1 (1:2,000, ThermoFisher MA5-17086), sGCβ1 (1:2,000, Cayman Chemicals,160897), AR (1:8,000, Santa Cruz Biotech, sc-816), GAPDH (1:10,000, Abcam, ab9485), cleaved-PARP (1:1,000, Cell Signaling, 9541), p16^INK4a^ (1:1,000, Abcam, ab108349), TRX1 (1:1,000, BD Biosciences, 559969),

HSP90 (1:2,000, BD Biosciences, 610418), phospho-VASP(Ser239) (1:1,000, Cell Signaling, 3114S), VASP (1:1,000, Santa Cruz, sc46668), β-actin (1:5,000, Abcam, ab8226), actin (1:2,500, Sigma, A2066), Bcl2 (1:2,000, Proteintech, 12789-1-AP), Survivin (1:1,000, Cell Signaling, 2808). Following incubation with the appropriate secondary horseradish peroxidase-conjugated antibodies, blots were developed using SuperSignal West Femto Maximum Sensitivity Substrate (ThermoFisher, 34095). Two molecular markers were used in this study: Cytiva Rainbow Molecular Weight Markers (Sigma, RPN800E) and Spectra Multicolor Broad Range Protein Ladder (ThermoFisher Scientific, 26634) depending on the size(s) of the protein(s) being targeted for detection. Densitometry of images was carried out via the ImageJ Analyze Gels (NIH) module and normalized to the loading signal for each band.

### Senescence-Associated Beta-Galactosidase Assay (SA-beta-gal)

SA-beta-gal staining was carried out as previously described^94^. Briefly, cells were washed in PBS, fixed in 0.2% glutaraldehyde for 5 min at room temperature, washed once in PBS and incubated overnight in freshly-made staining solution (1 mg/ml 5-bromo-4-chloro-3-indolyl-β-galactoside, 150 mM NaCl, 2 mM MgCl_2_, 5 mM K_3_Fe(CN)_6_, 5 mM K_4_Fe(CN)_6_, 40 mM NaPi, (pH 6.0). The next day, the cells were incubated for a further hour at 37 °C to intensify staining and then washed and stored in PBS at 4 °C till image acquisition. To quantify positive staining, 100 cells were counted for each sample over multiple fields of view, excluding fields at the very edge.

### Measurement of ROS

LNAI cells were plated into a 96-well black-walled plate at 2,000 cells/well in FBS culture medium and incubated at 37°C. Twenty-four hours after plating, cells were changed to their corresponding culture medium supplemented with 5% FBS or 5% CSS and treated with either 100 µM or 200 µM riociguat (Cayman chemical, 9000554) or DMSO (Sigma, D8418) as the vehicle control. At 20 h following riociguat treatment, cells were washed in ice-cold Ca^2^ and Mg^2+^ free 1X Hank’s balanced salt solution (HBSS, Thermo Fisher, 14175-095), and incubated with freshly prepared 10 µM dihydroethidium (hydroethidine) (HEt) (ThermoFisher, D11347), or 5- (and -6)-chloromethyl- 2′,7′-dichlorofluorescein diacetate (CM-H_2_DCFDA, ThermoFisher, C6827), or hydroxyphenyl fluorescein (HPF, ThermoFisher, H36004) for 30 min at 37°C. The cells were then washed in 1X HBSS once, and 150 μl 1X HBSS was added to each well on the plate prior to fluorescent detection. The plate was then read on the SpectraMax iD3 reader (Molecular Device) for detection of fluorescent signal. Unstained cells were used as negative controls. Data analysis and graphing were performed in GraphPad Prism (v.9).

### Single cell gel electrophoresis (comet) assay

Approximately 0.5 X 10^6^ LNAI cells were plated in 10cm dish in FBS media and incubated at 37°C. Twenty-four hours after plating, cells were changed into either media supplemented with 5% FBS or 5% CSS was added to the cells, and were treated with either 50 µM, 100 µM or 200 µM riociguat (Cayman chemical, 9000554) or DMSO vehicle control (Sigma, D8418). Drug and media were replenished daily for the total experimental duration (48 hours), after which the comet assay was carried out according to the comet assay kit (Trevigen, 4250-050-ES) instructions for alkaline unwinding and electrophoresis conditions. Gel electrophoresis was carried out at 21V for 30 min at 4 °C. Positive (CC0) and negative (CC3) control cells provided by Trevigen were run along with each sample to ensure that lack of tails or long tails in a sample were not due to technical issues. A minimum of 100 individual cells per sample were scored, with and without DNA tails.

### Xenograft tumor experiments

All animal studies were performed in accordance with the University of Miami Institutional Animal Care and Use Committee (IACUC)-approved protocol. Numbers of animals to be used for the tumor-formation experiments were determined through power analysis to provide 90% statistical power to detect a mean difference of 2.6 between two groups, assuming a two-sample Student’s t-test and a standard deviation for both groups of 1.5 at two-sided 5% significance level. For all xenograft experiments, cells were resuspended in a 1:1 matrigel (BD Biosciences, 356237): full FBS/RPMI-1640 media mixture and injected subcutaneously using a 26-gauge needle into one flank of immunocompromised 5–6-week-old castrated male mice (Nu/Nu, Envigo). Tumor length, width, and height were measured using electronic precision calipers (VWR, 90028).

Tumor volumes were calculated according to the following formula: 0.52x(heightxwidthxlength). For assessing whether riociguat could mitigate in vivo CRPC tumor growth, we subcutaneously injected 2 × 10^6^ LNAI cells. Animals were randomized into a vehicle or treatment groups once tumors were palpable (100–150 mm^3^) and treatment was initiated with either DMSO:Tween80:PBS (10%:10%:80%) or 20 mg kg^−1^ riociguat (Cayman Chemicals, 9000554) or cinaciguat (Cayman Chemicals, 17468) diluted in Tween80 and Mg^2+^ and Ca^2+^-free DPBS.

Injections were given intraperitoneally (IP) seven days a week starting at palpable tumor formation. For combinatorial ionizing radiation (IR)/riociguat studies, we subcutaneously injected 2 × 10^6^ LNAI cells on the flanks of 6-week-old, male, castrated Nu/Nu mice and allowed tumors to reach approximately 100 mm^3^ prior to randomization into vehicle and riociguat treated groups. After three days of daily IP treatment with either riociguat or vehicle, each group (vehicle and riociguat-treated) was randomized into either irradiated (6 Gy) or mock (0 Gy) groups. Non-tumor tissues were shielded with lead covers and tumors were subjected to one dose using the RadSource RS2000 as the radiation source. For assessing whether loss of the sGC heterodimer promoted castration resistance, we subcutaneously injected 2 x 10^6^ WT and KO sGCβ1 LNCaP cells into the flanks of 6-week-old, male, castrated Nu/Nu mice and allowed tumors to reach approximately 1000 mm^3^.

For all xenograft experiments, tumor measurements and animal weights were monitored three times a week in a non-blinded manner. Tumor-bearing animals were euthanized when tumors in any group exceeded 10% of animal body weight (∼1000 mm^3^). Immediately following euthanasia, blood was collected through cardiac puncture from experimental animals and processed for serum and plasma. Tumors were excised, cut sagittally where possible, photographed and sectioned into samples for formalin fixation or snap-frozen in liquid nitrogen. Lungs were also collected and snap frozen for systemic measurement of cGMP levels.

### Immunohistological stainining

For histopathological analysis, fine sections (4 μm) were cut from formalin-fixed, paraffin-wax-embedded samples, and stained with hematoxylin and eosin. Immunohistochemical analyses were performed utilizing ready-to-use Ki67-specific antibody solution (K2, Leica Biosystems, PA0230). Sections were dewaxed, rehydrated, and pretreated in a high pH (pH 9) solution at 100 °C. Slides were dehydrated using a standard protocol and mounted using Permount. Ki67 images were taken using a Leica microscope and associated LAS V4.9 software at 40X objective magnification.

For evaluation of tumor vasculature, thin-tissue FFPE slides were dewaxed and rehydrated as per standard protocols. Antigen retrieval was performed using low pH (pH 6) citrate solution at 15 PSI for 15 minutes. Following appropriate blocking PBS/10% goat serum, slides were incubated with primary antibody CD31 (Cell Signaling, CS77699) in PBS/1% BSA, followed by polymer treatment, 3% peroxide blocking, DAB chromogen (Vector, SK-4800), hematoxylin treatment (Leica, 3801), dehydrated and mounted using Permount. CD31-stained slides were imaged using the Olympus Whole Slide Scanner VS200 and processed using OlyVIA 2.9.1 software.

### Immunofluorescent staining

For determination of hypoxia, pimonadizole (Hypoxyprobe, HP10-100) was diluted in saline and administered to study animals intraperitoneally at a dose of 60 mg/kg two hours prior to euthanasia. Tissues (tumor, lung, and liver) were then collected and fixed in formalin and paraffin-embedded. Paraffin-embedded fine sections were deparaffinized by heating at 56°C for two hours and then hydrated by treating with Xylene (15 minutes, 2 times), 100% ethanol, 90% ethanol, and 70% ethanol (2 times) at 5 minutes each. The slides were steamed for 35 minutes with a pH 6 Dako Target Retrieval (Agilent Technologies, S169984-2) and permeabilized with 0.1% Triton X-100 (Sigma-Aldrich, T8787) for 5 minutes, followed by a 5-minute wash with PBS. Slides were then blocked in serum-free, protein blocker (Agilent Technology, X090930-2). Slides were incubated with a biotinylated anti-pimonidazole mouse IgG1 monoclonal antibody diluted 1:50 in Background Sniper (Fisher Scientific, 5082385) for one hour at room temperature. This step was followed by washes with PBS. Streptavidin, AlexaFluor 568 conjugated secondary antibody (Life Technologies, S11226) was applied for 30 minutes at a 1:200 dilution in the dark at room temperature followed by washes with PBS. Slides were then mounted using ProLong Gold anti-fade with DAPI (Molecular Probes, P36935) for immunofluorescence staining.

For CD44 immunofluorescence, the same procedure was followed except the primary antibody CD44 (Abcam, ab157107) was incubated on the slides at a 1:200 dilution in Background Sniper overnight at 4°C. Following this step, the secondary antibody, goat anti-rabbit IgG Superclonal Alexa Fluor 488 (Life Technologies, A27034) was diluted 1:200 in Background Sniper and applied for 30 minutes at room temperature.

Apoptosis was assessed by terminal deoxynucleotidyl transferase-mediated dUTP nick end-labeling (TUNEL), using the In Situ Cell Death Detection Kit, Fluorescein (Roche, 11 684 795 910). Slides were dewaxed, rehydrated and blocked as described and incubated with the TUNEL reaction mixture for 1 h at 37 °C in a humidified environment in the dark. After rinsing with PBS, the sections were mounted using Prolong Gold Antifade Mountant with DAPI. Images were acquired using a Leica fluorescence microscope.

### Cytokine Profiling

Sera from a vehicle-treated, riociguat-treated (non-responder; NR), and two riociguat-treated (responder; R) male mice bearing LNAI subcutaneous tumors were assayed for systemic cytokines using the Proteome Profiler Mouse XL Cytokine array (R&D Systems, ARY028). Membranes were exposed on film using the provided chemiluminescent reagent. Pixel densities of duplicate signals for the 111 cytokines were measured using the ImageJ Analyze Blots module.

### Measurement of cGMP levels

Steady state cGMP concentrations in cells, mouse plasma, tissues and human serum were measured by cGMP Enzyme Immunoassay (cat no. K065, Arbor Assays). Lysates were prepared according to the manufacturer’s instructions with minor modifications. Briefly, lung and tumor samples were pulverized in liquid nitrogen, 500 µl of sample diluent were added for every 50 mg of powdered lung tissue and 250 µl of sample diluent were added for every 50 mg of powdered tumor tissue. For measurement of cGMP levels in cell lines, cells were acutely treated with 20 µM riociguat or 20 µM cinaciguat and an equivalent DMSO dose for 30 minutes at 37 °C. The drug-containing media was removed, and cells were harvested by scraping in cold PBS and centrifuged at 3,500 rpm for 5 mins at 4 °C for for isolation of the cell pellet. Each pellet was resuspended in 150 μL sample diluent. All lysates were then sonicated and incubated on ice for 10 minutes for uniform lysis, followed by centrifugation at 13,000 rpm for 10 minutes at 4°C. The BCA assay was used to measure protein concentration of the lysates and cGMP levels were expressed as picomoles of cGMP per milligram of total protein for tissue samples. Measurements in sera and plasma samples were carried out as follows. Mouse plasma samples were diluted 1:5 in sample diluent. Human sera from prostate cancer patients were assayed undiluted using the Cyclic GMP Direct Assay, as cGMP is lower in sera than plasma. An acetylation format was used to allow low concentrations of cGMP to be detected in tissue, cells, serum, and plasma. The cGMP signal was read and calculated at 450 nm on a Spectramax iD3 or iDx Microplate Reader. Plasma and serum cGMP level were multiplied by their respective dilution factors and expressed in nM concentration.

### Determination of PSA levels

Serum was collected through centrifugation of blood samples and diluted 1:2 in PBS. The PSA ELISA assay was performed for each serum sample in triplicate, as per manufacturer’s instructions for the PSA ELISA kit (Biocheck, BC-1019). Absorbances were read at 450 nm on a Spectramax iD3 Microplate Reader. Final values were in ng ml^-1^ and reflected adjustments based on initial dilution of serum where necessary.

### Statistical Analyses

Specific descriptions of statistical analyses and experimental/biological replicates are provided in the figure legends and in relevant Methods subsections. Data are presented as ± standard error of the mean (SEM). Data were analyzed by two-tailed unpaired Student’s *t*-test, analysis of variance (ANOVA) for group analyses or Spearman correlation tests. All tests for comparing area under the curve (aAUC) used the Wilcoxon test for two groups, or the Kruskal-Wallis test for more than two groups. Dose response curves were estimated using 4-parameter log-logistic function and the IC_50_ was obtained from the estimated parameters. Results with p-values < 0.05 were considered statistically significant, with * p< 0.05, ** p≤ 0.01, *** p≤ 0.001, **** p≤ 0.0001. Statistical analyses were performed using statistical software package R (v. 4.2.2) or GraphPad Prism (v. 9).

## Supporting information

all supplementary data

## Acknowledgments

This work was supported by R01CA254100 and W81XWH-16-1-0643 (PR), R01GM067640 and R01GM112415 (AB), F31CA232653 (CT), Paul F. Calabresi K12 (JS), a Sylvester Translational Science Grant (PR, ACL), a UMMSOM Summer Undergraduate Research Fellowship (LML), and research support funds from the Sylvester Comprehensive Cancer Center (PR). We thank Dr. Roger Alvarez for helpful discussions, and Mr. Ayush Rana and Mr. Philip Gregory for assistance with experiments. We thank Dr. Jenn Marte at the NCI. This study utilized the following Sylvester Shared Resources: Bioinformatics and Biostatistics Shared Resource, Onco-Genomics Shared Resource, and Cancer Modeling Shared Resource. Research reported in this publication was supported by the National Cancer Institute of the National Institutes of Health under Award Number P30CA240139. The content is solely the responsibility of the authors and does not necessarily represent the official views of the National Institutes of Health.

