## Supplementary material for "The soluble guanylyl cyclase pathway is inhibited to evade androgen deprivation-induced senescence and enable progression to castration resistance": all supplementary data

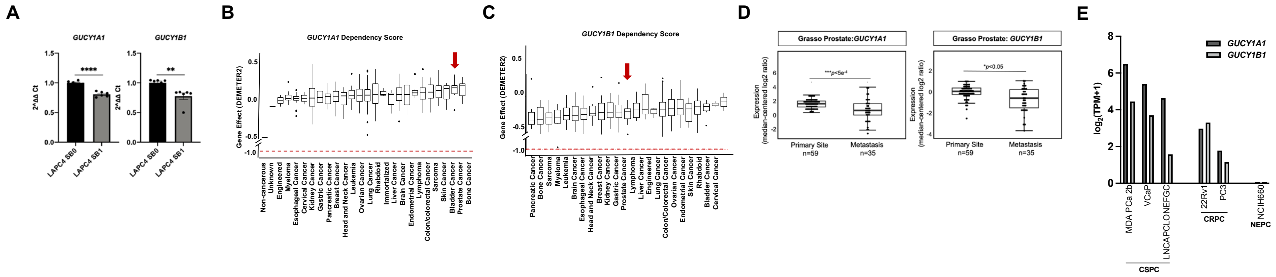


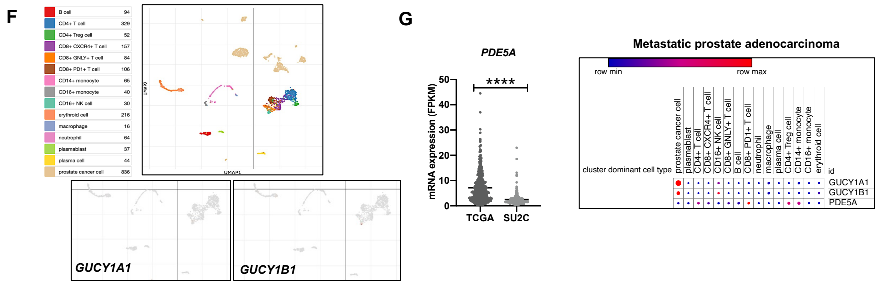


H

| Patient | Age | Gleason | Race | mCSPC (months) | mCRPC (months) |
|---|---|---|---|---|---|
| 25 | 45 | 9 | White | 10.5 | 35.5 |
| 35 | 86 | 6 | Black | 115 | 31.5 |
| 42 | 50 | 9 | Black | 10.5 | 42 |
| 45 | 55 | 9 | White | 12.5 | 16 |
| 48 | 57 | 9 | White | 26 | 29 |
| 49 | 73 | 9 | Black | 14.5 | 40 |
| 61 | 78 | 9 | Hispanic | 19 | 27 |
| 62 | 66 | 8 | White | 37.5 | 31 |
| 64 | 67 | 9 | White | 21.5 | 24.5 |
| 71 | 63 | 9 | Hispanic | 36 | 8 |

**Supplementary Figure S1**. **Cancer cells are not dependent on sGC expression for viability**

Note that all error bars represent ± SEM and * p< 0.05, ** p≤ 0.01, *** p≤ 0.001, **** p≤ 0.0001.

A. mRNA expression of *GUCY1A1*, *GUCY1B* in LAPC4 SB0 and LAPC4 SB1 were analyzed by qPCR. *ActinB* was used for normalization. Unpaired two-tailed Student's t tests were used to determine p-values.

B. *GUCY1A1* dependency profiles of indicated cancer types from DepMap.org. The red line shows the gene effect threshold below which a cancer line is deemed dependent on a particular gene. Prostate cancer is indicated by a red arrow.

C. *GUCY1B1* dependency profiles of indicated cancer types from DepMap.org with the same metrics denotes as in (B).

D. Box-and-whisker plots showing *GUCY1A1 and GUCY1B1* mRNA expression. Data were derived from the Grasso primary vs. metastatic datasets[17]. The datasets represent primary vs metastatic human prostate tumors. Boxplots represent the five-number distribution with the top and bottom of the boxes indicating the 75th and 25th percentile, respectively. The whisker represents 1.5 times the interquartile range from the box. Number of samples (n) and p-values (determined by a two-tailed Mann–Whitney *U* test) are as shown.

E. *GUCY1A1* and *GUCY1B1* mRNA expression levels for the indicated cell lines mined from DepMap.org.

F. The full UMAP plot for all clustered populations from the metastatic treatment resistant tumors depicted in Fig. 1F (top). Expression of *GUCY1A1* and *GUCY1B1* in tumor immune cells (bottom).

G. Left: *PDE5A* mRNA levels from the TCGA vs SU2C prostate cancer dataset (cbioportal.org). The p-values were determined via an unpaired two-tailed Student's t test. Right: Analysis of *GUCY1A1*, *GUCY1B1* and *PDE5A* expression in cancer and indicated immune cells from the human metastatic prostate cancer in Fig. 1F[19].

H. Characteristics and clinical outcomes of patients whose sera were profiled for cGMP (Fig1I, J).

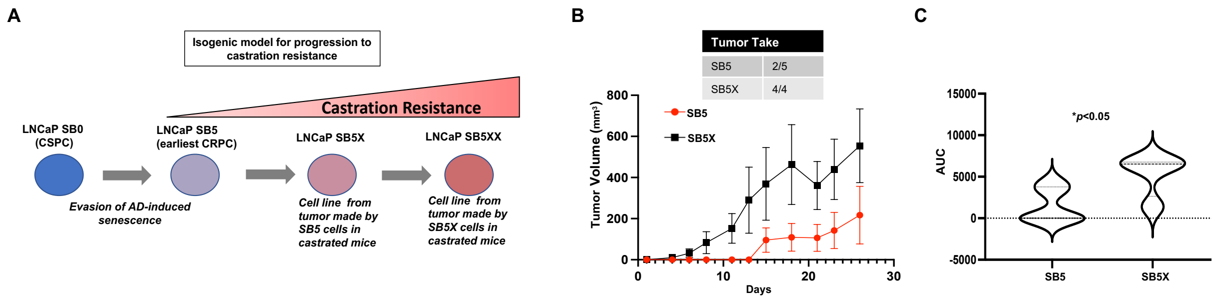


**Supplementary Figure S2. LNCaP SB0-derived emergent and progressively castration-resistant variants**

A. Schematic showing development of isogenic LNCaP-derived lines representing our model of progressive castration resistance.
B. Subcutaneous xenograft tumors were established from LNCaP SB0 and LNCaP SB5 cells. Tumor growth kinetics and tumor incidence per injection site are shown.
C. Violin plots representing the tumor growth curve AUCs from (B). The p values compare aAUC of the curves using a Wilcoxon two-sample test, with $p < 0.05$ considered significant.

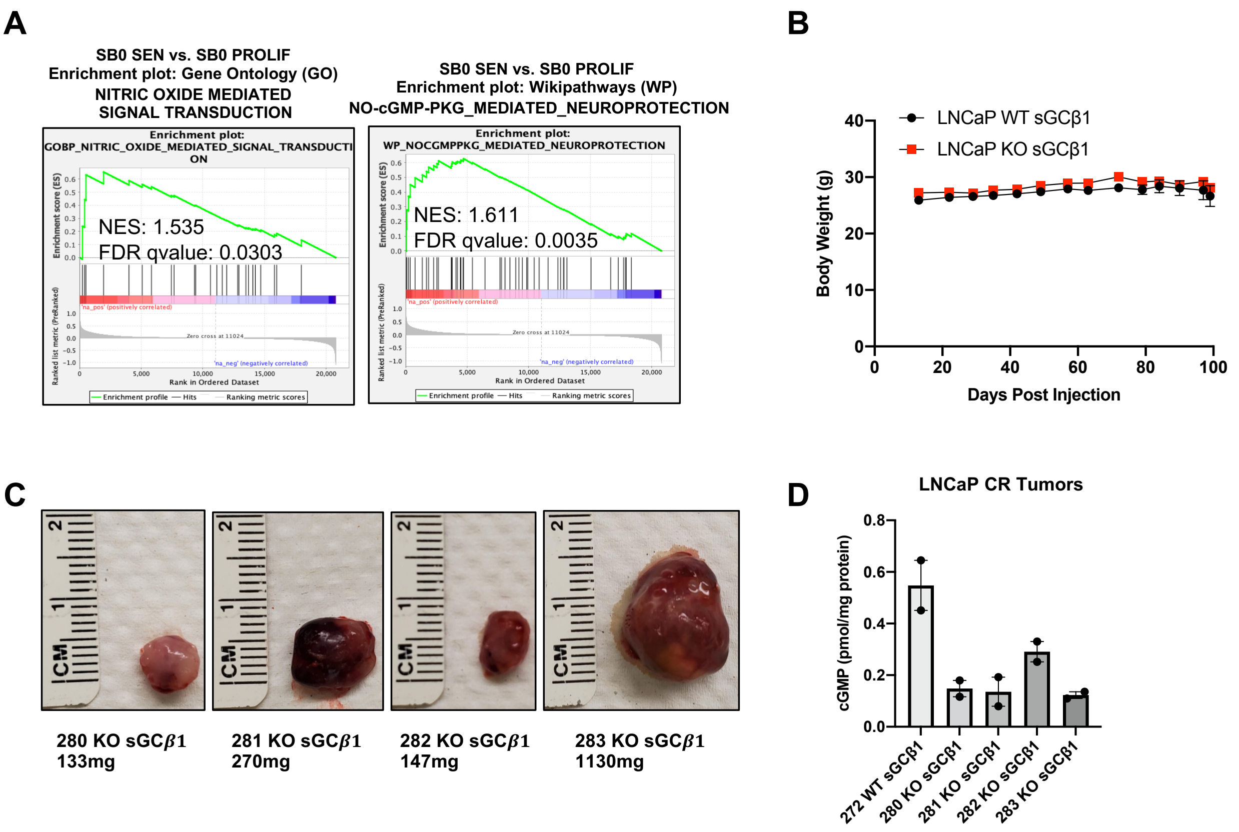


**Supplementary Figure S3. Deletion of the sGCβ1 subunit in LNCaP CSPC cells facilitates castration resistance**

A. Gene set enrichment analysis of our Illumina data demonstrated a positive enrichment of two Gene Ontology (GO) pathways associated the NO-sGC-cGMP signaling in the ADIS induction dataset from Fig. 1B. Normalized enrichment score (NES) and FDR-adjusted p-values are shown.
B. Animal body weights from the experimental xenograft tumor groups described in Fig. 3I are shown. Error bars represent ± SEM.
C. Representative images of subcutaneous LNCaP KO sGCβ1 (n=4) tumors formed in castrated male Nu/Nu mice. Tumors were weighed immediately following resection and the weights are noted below the images. The sole wildtype tumor was not able to wholly resected due to its small size and therefore could not be photographed; however small sections were collected and flash-frozen for cGMP and protein lysates.
D. Intratumoral cGMP levels from the sole WT sGCβ1 and four KO sGCβ1 tumors. Error bars represent ± SEM.

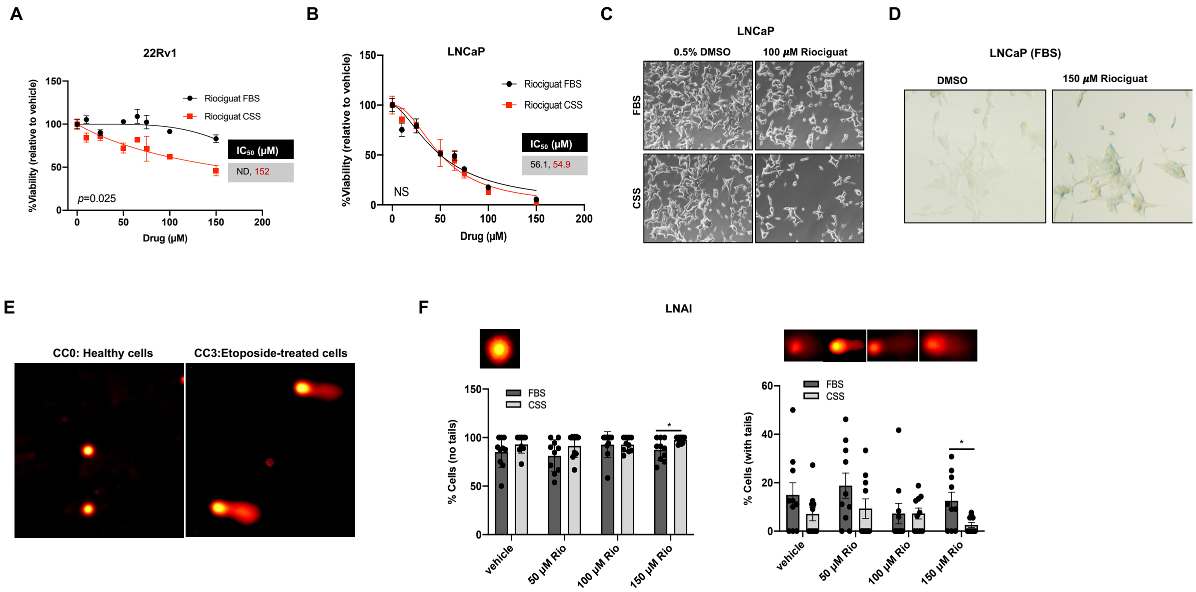


**Supplementary Figure S4. AD-induced modulation of riociguat activity and its anti-tumor effects in CRPC vs. CSPC cell lines**

Note that all error bars represent ± SEM and * p< 0.05, ** p≤ 0.01, *** p≤ 0.001, **** p≤ 0.0001.

A. 22Rv1 cells were plated in 96-well plates in triplicate and treated with vehicle (DMSO) or riociguat in FBS- or CSS-supplemented media for 72 hours with daily redosing. Cell viabilities, $IC_{50}$s and p-values were established as in Fig. 4C.

B. LNCaP cells were plated at 96-well plates in triplicate and treated with riociguat in FBS- or CSS-supplemented media for 72 hours with daily redosing. Cell viabilities, $IC_{50}$s and p-values were established as in Fig. 4C.

C. LNCaP cells were plated and were treated with vehicle (DMSO) or riociguat in FBS- or CSS-supplemented media for 72 hours with daily redosing. Light microscopy images of cells were taken at 10X.

D. LNCaP cells were plated and treated with vehicle (DMSO) or riociguat in FBS-supplemented media for 72 hours with daily redosing. Light microscopy images of cells were taken at 10X, following the SA-beta-gal assay.

E. Representative cell images for negative (CC0) and positive (CC3, etoposide treated) technical controls (Trevigen) for the comet assay.

F. LNAI cells were plated in 10cm dishes and treated with vehicle (DMSO) or riociguat in either FBS- or CSS-supplemented media. The comet assay was performed following a 48-hour treatment, with daily riociguat redosing. A minimum of 100 individual cells per condition were scored and p-values were established using an unpaired two-tailed Student's t test, with p < 0.05 considered significant. Representative cell images are shown above the respective graphs.

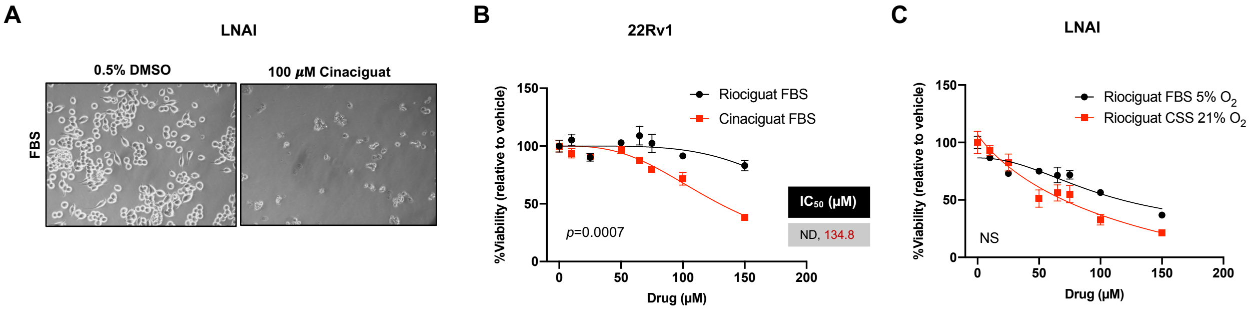


**Supplementary Figure S5. Response of CRPC cells to the sGC activator, cinaciguat**

Note that all error bars represent ± SEM and * $p < 0.05$, ** $p \leq 0.01$, *** $p \leq 0.001$, **** $p \leq 0.0001$.

A. LNAI cells were treated with vehicle (DMSO) or cinaciguat in FBS-supplemented media for 72 hours with daily redosing. Light microscopy images of cells were taken at 10X.
B. 22Rv1 cells were plated at 96-well plates in triplicate and treated with riociguat or cinaciguat in FBS-supplemented media for 72 hours with daily redosing. Cell viabilities, $IC_{50}$s and p-values were established as in Fig. 4C.
C. LNAI cells were plated at 96-well plates in triplicate and treated with riociguat in FBS-supplemented media at 5% $O_2$ or in CSS-supplemented media at 21% $O_2$ for 72 hours with daily redosing, to compare relative sensitivities to riociguat. Cell viabilities, $IC_{50}$s and p-values were established as in Fig. 4C.

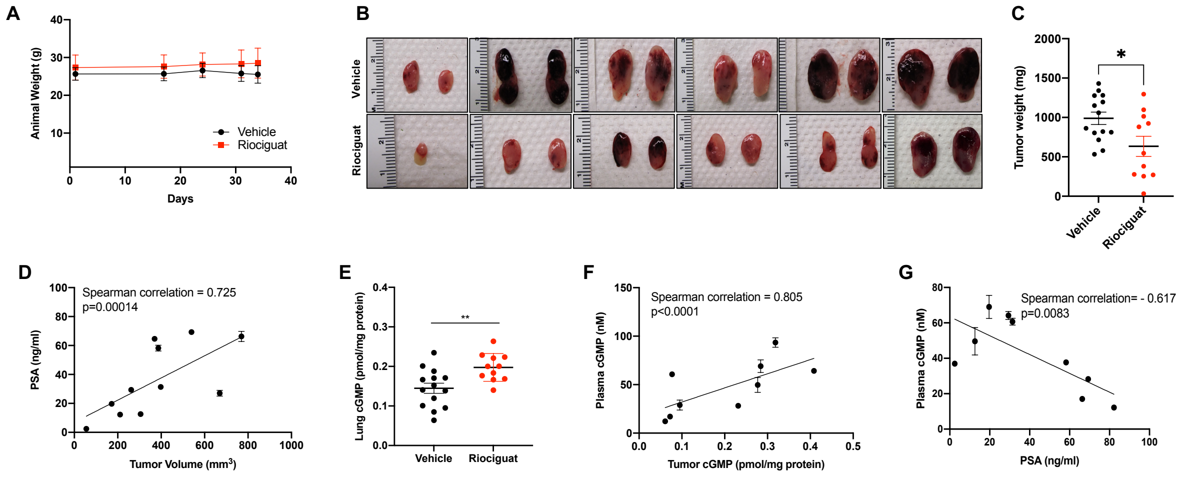


**Supplementary Figure 6**. **Riociguat-treated LNAI CRPC resected tumors, tumor weights, PSA levels and cGMP levels**
Note that all error bars represent ± SEM and * $p < 0.05$, ** $p \leq 0.01$, *** $p \leq 0.001$, **** $p \leq 0.0001$.

A. Mean body weights per treatment group (from Fig. 6A) at the indicated time points in the tumor growth curve are shown.
B. Representative images of castration-resistant LNAI xenograft tumors derived from vehicle-treated and riociguat-treated groups in Fig. 6.
C. Endpoint tumor weights (mg) of castration-resistant LNAI xenograft tumors derived from the vehicle-treated and riociguat-treated group (Fig. 6A). Tumors were weighed immediately post-resection. Two-tailed Student's t-test with Welch's correction was used to determine p-values.
D. PSA levels in riociguat-treated tumors correlate positively with tumor volume. Each point represents PSA values from an individual tumor. Spearman correlation test values are shown.
E. Pulmonary cGMP levels were measured in lung lysates from the vehicle or riociguat-treated groups in Fig. 6A. Unpaired two-tailed Student's t-test was used to determine p-values.
F. Plasma cGMP levels from riociguat-treated animals correlate positively with intratumoral cGMP levels. Spearman correlation test values are shown.
G. PSA levels from riociguat-treated animals correlate inversely with their plasma cGMP levels. Spearman correlation test values are shown.

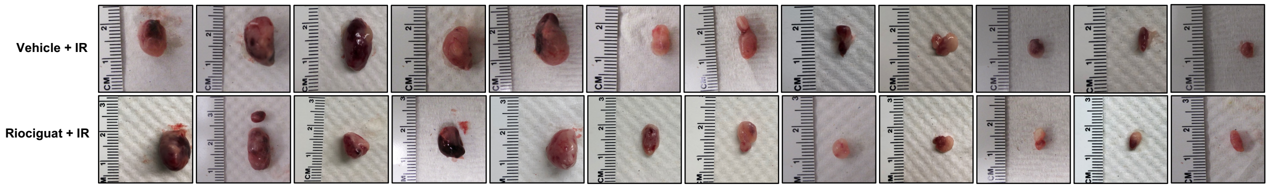


**Supplementary Figure 7. Representative resected tumor from riociguat/IR treated animals**
Representative images of subcutaneous tumors derived from the vehicle + IR and riociguat + IR groups (Fig. 7E).
